# Laminar-specific control of response gain and orientation-tuning by parvalbumin-expressing inhibitory interneurons in primate visual cortex

**DOI:** 10.64898/2025.12.23.696300

**Authors:** Amin Vafaei, Andrew M. Clark, Alireza Khadir, Weifeng Dai, Frederick Federer, Alessandra Angelucci

## Abstract

Understanding the role of the different types of inhibitory interneurons in cortical computations is central to elucidating how the neocortex processes sensory information. The emergence of orientation tuning in primate primary visual cortex (V1) is a canonical model for studying how cortical sensory circuits and inhibitory interneurons compute relevant stimulus features. The selective feedforward convergence of non–orientation-selective thalamic afferents establishes initial orientation tuning in the granular V1 input layer. As signals propagate through the cortical microcircuit, orientation tuning sharpens in extra-granular layers, yet the underlying mechanisms and the contribution of specific inhibitory neuron subtypes within V1 remain unresolved. To study the role of the largest cortical inhibitory neuron subclass, parvalbumin-expressing (*PV*^+^) interneurons, in this V1 computation, we combined laminar extracellular recordings with bidirectional optogenetic manipulations of *PV*^+^ cells in marmoset V1. Our results reveal a striking laminar specificity: in granular layers, *PV*^+^ cells implement divisive/ multiplicative linear gain control, whereas in extra-granular layers they exert tuned nonlinear suppression that enhances orientation tuning. Computational modeling suggests that *PV*^*+*^ neurons can control gain by changing a neuron’s spiking threshold, and orientation tuning by changing a neuron’s input noise, which regulates the neuron’s input–output function. Our findings reconcile divergent results from previous rodent studies and establish a framework for understanding layer-dependent inhibitory computations in the primate cortex.

## INTRODUCTION

Understanding the role of inhibitory interneurons in shaping cortical activity is fundamental to elucidating how the brain processes sensory information. Inhibitory interneurons, though a minority of cortical cells, are indispensable for shaping cortical computations, regulating the excitatory drive and sensory responses of pyramidal (Pyr) neurons through mechanisms that control gain, suppress noise, and coordinate spike timing^1–5^. Among the major subclasses, parvalbumin-expressing (*PV*^+^) interneurons—comprising fast-spiking basket and chandelier cells^6,7^ — are particularly well-positioned to influence cortical computations due to their intrinsic properties and connectivity patterns. Specifically, *PV*^+^ cells constitute the most prevalent subclass of cortical inhibitory interneurons and are characterized by fast-spiking activity, non-adapting firing patterns, and axonal projections targeting the soma, perisomatic compartments, and axon initial segment of Pyr neurons, thus enabling rapid and potent Pyr neuron inhibition^8-10^. Despite their ubiquity, whether the role of *PV*^+^ cells in sensory processing is uniform across cortical layers and conserved across species remain important open questions.

The well-studied phenomenon of orientation tuning in primary visual cortex (V1) provides an attractive model for investigating the role of inhibitory interneurons in cortical computations. Since its discovery in cat V1^11^, how the cortex computes orientation tuning from untuned thalamic inputs, has been a highly debated question in neuroscience. In particular, whether intracortical inhibition is required to generate tuning, or whether the latter can arise from the intrinsic non-linearities of thalamic afferents and cortical neurons, remains a central question^12-14^. This question, and in particular the role of distinct inhibitory neuron types in orientation tuning, has been extensively investigated in the mouse, where cell-type specific optogenetics became feasible much earlier than in higher mammals.

Despite extensive investigation, the contribution of *PV*^+^ neurons to orientation selectivity in mouse V1 remains contested. One study reported that *PV*^*+*^ neuron activation produces subtractive suppression and sharpens orientation tuning^15^, while other studies reported divisive/multiplicative effects of *PV*^*+*^ neuron manipulation without changes in tuning sharpness^16,17^. Much of the discrepancy across studies was attributed to differences in optogenetic activation parameters^18^, as weak or brief *PV*^+^ activation often left tuning unchanged, whereas strong, sustained activation produced robust sharpening. These observations suggest that *PV*^+^ effects on orientation tuning may be strongly context-dependent, potentially varying with cortical layer, input–output balance, and the temporal dynamics of activation.

Another important consideration is that orientation tuning emerges differently across species. In rodents, orientation selectivity is present prior to V1, in both the retina and lateral geniculate nucleus (LGN) of the thalamus^19-23^; moreover, orientation-tuned cells in V1 receive selective inputs from orientation-tuned cells in LGN and retina, suggesting their tuning may be largely generated subcortically^24^. In contrast, in primates and carnivores, orientation tuning largely appears in V1, and, in primates, is weak in the input, granular, layer of V1 and is sharpened outside this layer^25-27^. These, and other interspecies differences, such as the relative prevalence and different laminar distribution of *PV*^+^ cells^28-32^ raise the question of whether mechanisms underlying orientation tuning—and the role of inhibitory neurons in shaping it—are conserved or divergent across taxa.

Here we have examined the role of *PV*^+^ cells in orientation tuning in marmoset V1, by combining laminar-resolved extracellular recordings with bidirectional optogenetic manipulation of *PV*^+^ interneurons over a wide range of manipulation strengths. Using drifting gratings and precise laminar classification, we show that *PV*^+^ neuron activation/inactivation produces linear divisive/multiplicative scaling in granular layers—consistent with gain control—and nonlinear (tuned or Mexican-hat-like) changes in extra-granular layers, which sharpen/broaden orientation tuning. Using computational modeling, we show that this nonlinear effect is described by the noise-controlling function of *PV*^+^ cells in extra-granular layers, which regulate the input–output function of Pyr cells. These laminar-specific effects reconcile divergent findings from rodent studies and point to important inter-species differences, by demonstrating that *PV*^+^ neuron function depends critically on cortical layer, establishing a framework for understanding inhibitory circuit computations in the primate visual cortex.

## RESULTS

To examine the role of V1 *PV*^+^ interneurons, we selectively manipulated *PV*^+^ cells optogenetically while monitoring their impact on Pyr neuron visual responses. For bidirectional control of *PV*^+^ activity, we selectively expressed in *PV*^+^ cells, in V1 of two separate hemispheres, either the light-sensitive cation-conducting channelrhodopsin *C1V1*^33^, for *PV*^+^ activation (PVA), or the soma-targeted anion-conducting channelrhodopsin *stGtACR2*^34^, for *PV*^+^ inactivation (PVI) (**Fig. 1A**). The two opsins were delivered via injections of the viral vector AAV-PhP.eB under the control of the *S5E2* enhancer^35^, which selectively and efficiently transduces *PV*^+^ cells in marmoset V1^31^ (see STAR Methods). After 4-6 weeks, we optogenetically manipulated *PV*^+^ neurons activity by surface photostimulation of varying intensity, while recording extracellular neural activity using high-density (24- or 64-channels) laminar electrode arrays (LEAs) oriented perpendicular to the surface of V1, and targeted to a region of opsin expression. Here we present data from a total of 11 LEA penetrations (669 spike-sorted single units, 302 for PVI and 367 for PVA) recorded in 6 hemispheres of 4 sufentanil-anesthetized marmosets (see STAR Methods).

### *PV*^+^ Interneuron Manipulation Exerts Distinct Effects on Pyr Neuron Responses

We recorded neuronal responses through a V1 column while presenting drifting grating stimuli of optimal parameters for most neurons in the column varying in direction, to measure orientation tuning functions under control (no laser) and optogenetic manipulation (laser) conditions (**Fig. 1 B-C**). Trial interleaved, focal surface laser photostimulation of varying intensity was applied at the V1 recording site throughout the duration of visual stimulus presentation (1 s with 2 s interstimulus interval; see STAR Methods). We analyzed data from a total of 537 orientation-selective single units (310 for PVA and 227 for PVI).

PVA reduced the stimulus-evoked firing rate in 90% (278 of 310) of recorded orientation-selective units, whereas PVI increased it in 95% (217 of 227) of units. The total fraction of cells (8%) exhibiting increases (decreases) in firing rate in response to PVA (PVI), considered putative *PV*^+^ cells, is consistent with the estimated prevalence of *PV*^+^ cells (~10%) in marmoset V1 and the efficiency of the *S5E2* enhancer in transducing *PV*^*+*^ cells^31^. These putative *PV*^+^ cells (n=42) were excluded from further analyses. Therefore, we analyzed data for a total of 495 orientation-selective non-*PV*^*+*^cells.

Previous optogenetic studies in mouse V1 have reported three principal effects of *PV*^*+*^ cell manipulation on Pyr cells’ orientation-tuning functions: subtractive shifts^15^, divisive scaling^16^, or a combination of both^17^. Moreover, all of these effects could be explained by a threshold linear model with both multiplicative/divisive and additive/subtractive components^36^. We asked whether we could identify comparable effects in our data. **Fig. 1B-C** shows visual responses and orientation-tuning functions for 4 representative single units recorded under control and laser conditions for PVA and PVI. As expected, PVA suppressed neuronal firing rates, while PVI increased it. Consistent with previous reports^15,17^, *PV*^+^ manipulation did not alter the preferred orientation of these cells, nor across the population (**Fig. S1A**), but changed tuning functions in different ways. To quantify these changes, we computed two measures. The first measure (Δ*T* = *T*_lowFr_ − *T*_highFr_) is the difference between the tuning curve with lower firing rates and the tuning curve with higher firing rates. Thus, *T*_lowFr_ corresponds to the laser condition in PVA, but the control condition in PVI (both corresponding to conditions of higher *PV*^*+*^ neuron activity). Δ*T* is expected to yield a flat horizontal line if the inhibitory effect of *PV*^*+*^ interneurons is linear additive/subtractive. The second measure calculates the ratio between the tuning curves in the two conditions, 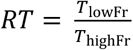. *RT* is expected to yield a flat horizontal line if the inhibitory effect of *PV*^*+*^ interneurons is linear divisive/ multiplicative. For the two units in **Fig. 1B**, *RT* yielded a flat line (*cyan curve*), while Δ*T* yielded a difference curve with the strongest effect of PVA/PVI at the recorded neuron’s preferred orientation (*red curve*). Thus, for these two units the effect of PVA/PVI was linear divisive/multiplicative (D/M effect). In contrast, for the two units in **Fig. 1C** the difference curve (Δ*T*) had a Mexican-hat profile, i.e. stronger effect of the optogenetic manipulation at orientations away from the preferred (in both cells strongest at the oblique orientations), and the ratio curve (*RT*) had a non-flat profile, with weakest effects at the preferred orientation, indicative of non-linear transformations (NL effect). The bottom row of **Fig. 1B-C** plots the response of each example unit to gratings of each orientation in control vs. laser conditions (normalized to the max response in the control condition), fitted with the following threshold linear function, as in Atallah et al.^17^ (see STAR Methods):

**Figure 1.**
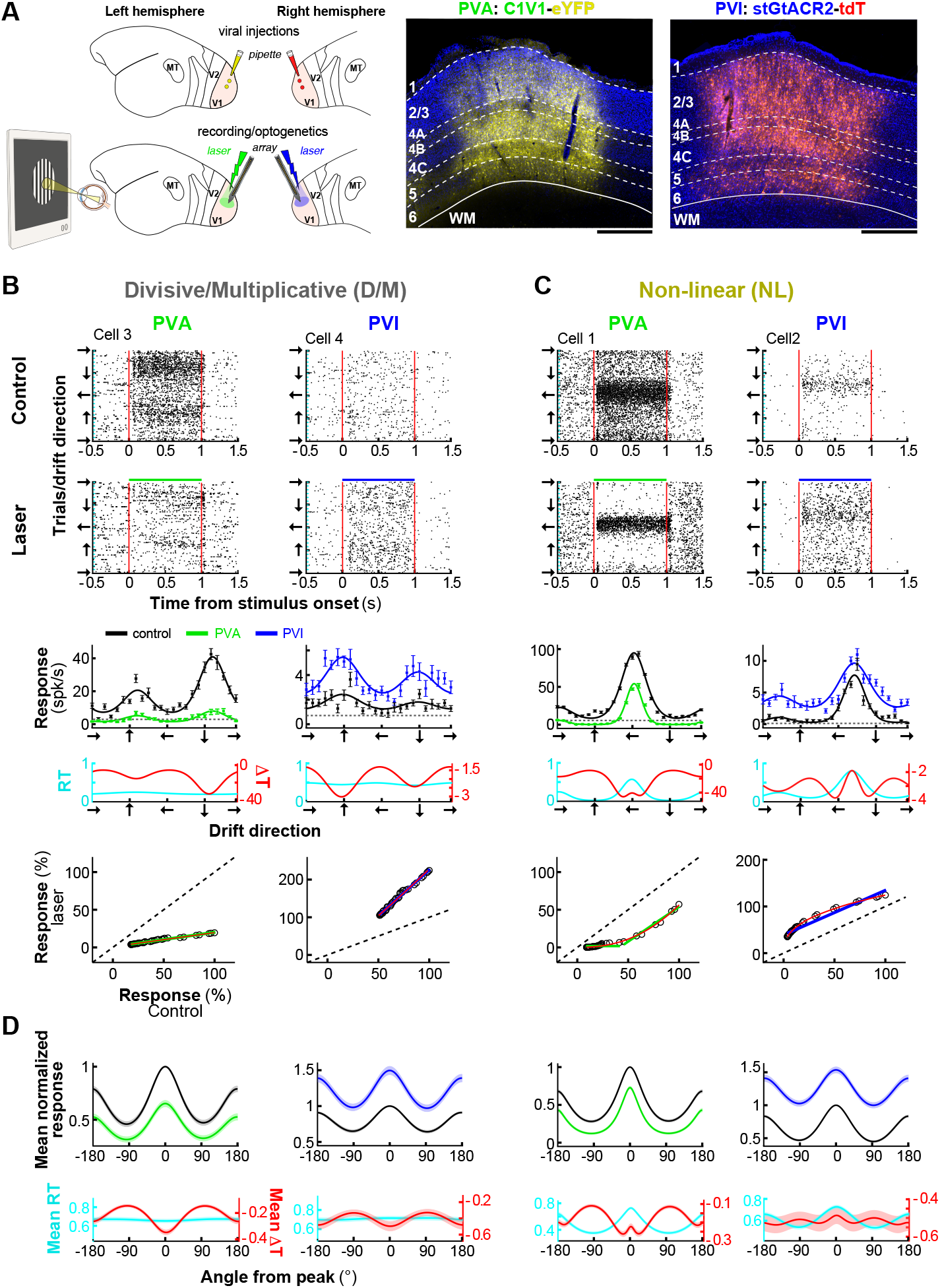
Distinct effects of *PV*^+^ interneuron manipulation on Pyr neurons’ responses. **(A)** <u>Left:</u> schematics of the experimental paradigm. <u>Top Left:</u> on day 1 a viral vector carrying the genes for C1V1 and eYFP is injected in one hemisphere, and a vector carrying the genes for stGtACR2 and tdT is injected in the contralateral hemisphere. <u>Bottom Left</u>: 4-6 weeks later, a terminal optogenetic/recording experiment is performed. <u>Middle and Right:</u> micrographs of V1 tissue sections across V1 layers (layer borders marked by *white dashed contours*) showing expression of C1V1-eYFP (*yellow*) and stGtACR2-tdT (*red*) in *PV*^*+*^ cells, used for *PV*^*+*^ cell activation (PVA) and inactivation (PVI), respectively. Scale bars: 0.5 mm. **(B)** <u>First and second row:</u> Raster plots (Y axis indicates individual trials for stimuli of different drift direction) for two example single units under control (<u>First row</u>) and laser (<u>Second row</u>) conditions, for PVA (<u>Left</u>) and PVI (<u>Right).</u> Visual stimuli were presented for 1s and photostimulation was simultaneous with visual stimulus presentation (second row, *green and blue bars* at the top of the rasters). *Red vertical lines* mark stimulus onset and offset. <u>Third row</u>: orientation tuning curves in control (*black*) and laser (PVA *green*, PVI *blue*) conditions fitted with von Mises functions (see STAR Methods). Error bars: s.e.m. *Black dashed line*: baseline activity. <u>Fourth row</u>: *ΔT* (*red*) shows the difference between the tuning function with lower firing rate and the function with higher firing rate. *RT* (*cyan*) shows the ratio between the same two tuning functions. <u>Fifth row</u>: Spiking responses (% of max response in control condition) of the same units to stimuli of different drift directions in control vs laser conditions for PVA (<u>Left)</u> and PVI (<u>Right</u>). G*reen and blue lines*: threshold linear model fits. *Red line:* fits of our I/O model described later in the Results. The example units in (B) showed divisive/multiplicative (D/M) effects of *PV*^*+*^ cell manipulations. **(C)** Same as in (B) but for two example units showing non-linear (NL) effects of *PV*^*+*^ cell manipulations. **(D)** <u>Top:</u> Normalized population-averaged tuning curves ± s.e.m. (*shading*) for units grouped by effect type (D/M: Left two columns; NL: right two columns) and manipulation direction (PVA: first and third column; PVI: second and fourth column). <u>Bottom:</u> averaged *RT* (*cyan*) and *ΔT* (*red*) curves for the populations grouped by effect type and manipulation direction. The method used for clustering units by effect type is shown in **Fig. S2. Fig. S3** shows examples of *PV*^*+*^ neuron manipulation effects on Unclassified Pyr neuron responses.

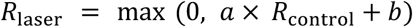

Where, *R*_laser_ and *R*_control_ are the response in the laser and control condition, respectively. According to this model, the optogenetic effect of *PV*^*+*^ cell manipulation is linear and should, thus, largely preserve the shape of the tuning function (D/M changes above zero firing rate), and Δ*T* curves should be proportional to the control tuning functions. As expected, this model fitted D/M units well (**Fig. 1B**), but not the NL units (**Fig. 1C**).

Based on Δ*T* and *RT* curve profiles, we classified all orientation-selective units in our recorded sample (n=495) into three groups: NL, D/M, and Unclassified (**Fig. S2**, see STAR Methods). The majority of recorded units (54%, n=266/495) exhibited NL effects (PVA: n=131/266 units; PVI: n=135/266); a smaller proportion (29%, n=146/495) showed D/M effects (PVA: n=88/146; PVI: n=58/146), and 17% of units (n= 83/495) could not be classified as NL or D/M (Unclassified, U). The U cells included a small group of cells (Mix, n=29) which showed both NL and D/M effects at different laser intensities. In contrast to these cells, the cells classified as NL and D/M showed the same effect at all laser intensities, as verified by matching spike-sorted waveforms across laser intensities. The remaining U cells (uncategorized-Uct, n=54 units; **Fig. S2B**) showed neither NL or D/M effects; this group included cells for which PVA abolished responses at all orientations (e.g. cells 5,8 in **Fig. S3**), cells with noisy manipulation effects (e.g. cell 7 in **Fig. S3**), and cells that did not show NL or D/M effects (e.g. cell 6 in **Fig. S3**). Notably, we found no purely additive/subtractive effects (i.e., no flat Δ*T* curves, **Fig. S2C**). To quantify the “flatness” of the Δ*T* curve, we measured the Coefficient of Variation (CoV) as the standard deviation (SD)/mean of the Δ*T* curve. Both the D/M and NL populations showed large CoV values (mean±s.e.m, D/M: 0.432±0.0183, PVA; 0.251±0.0253, PVI; NL: 0.377±0.0148, PVA; 0.177±0.0137, PVI) indicative of lack of subtractive/additive effects of *PV*^*+*^ cell manipulations (**Fig. S2D**; see also STAR Methods *Neuronal sample selection and clustering*).

**Fig. 1D** shows the population-averaged normalized tuning curves for D/M and NL units grouped by effect type, in control and laser condition (pooled across laser intensities), and the average Δ*T* and *RT* curves for each population (Δ*T* and *RT* curves for all individual units are shown in **Fig. S2C**). These population averaged tuning curves showed similar behavior as described for the representative single units in **Fig. 1B-C**.

### Impact of *PV*^+^ Interneurons on Pyr Neuron Orientation Tuning

We next asked how *PV*^+^ interneurons influence Pyr neurons’ orientation tuning and selectivity. For the D/M and NL populations, separately, **Fig. 2A,E** plots the change in half-bandwidth (ΔHBW= HBW_laser_-HBW_control_), a metrics of tuning sharpness (see STAR Methods), caused by the optogenetic manipulation, as a function of percent change in firing rate (ΔFr), caused by photostimulation at varying irradiance. We used as a proxy of laser intensity the laser-induced percent change in firing rate rather than irradiance; this allowed us to compare effects across penetrations, injection sites and cells at different depths, as a given irradiance can cause different effect magnitudes depending on opsin expression levels and cell distance from the light source (see also STAR Methods).

**Figure 2.**
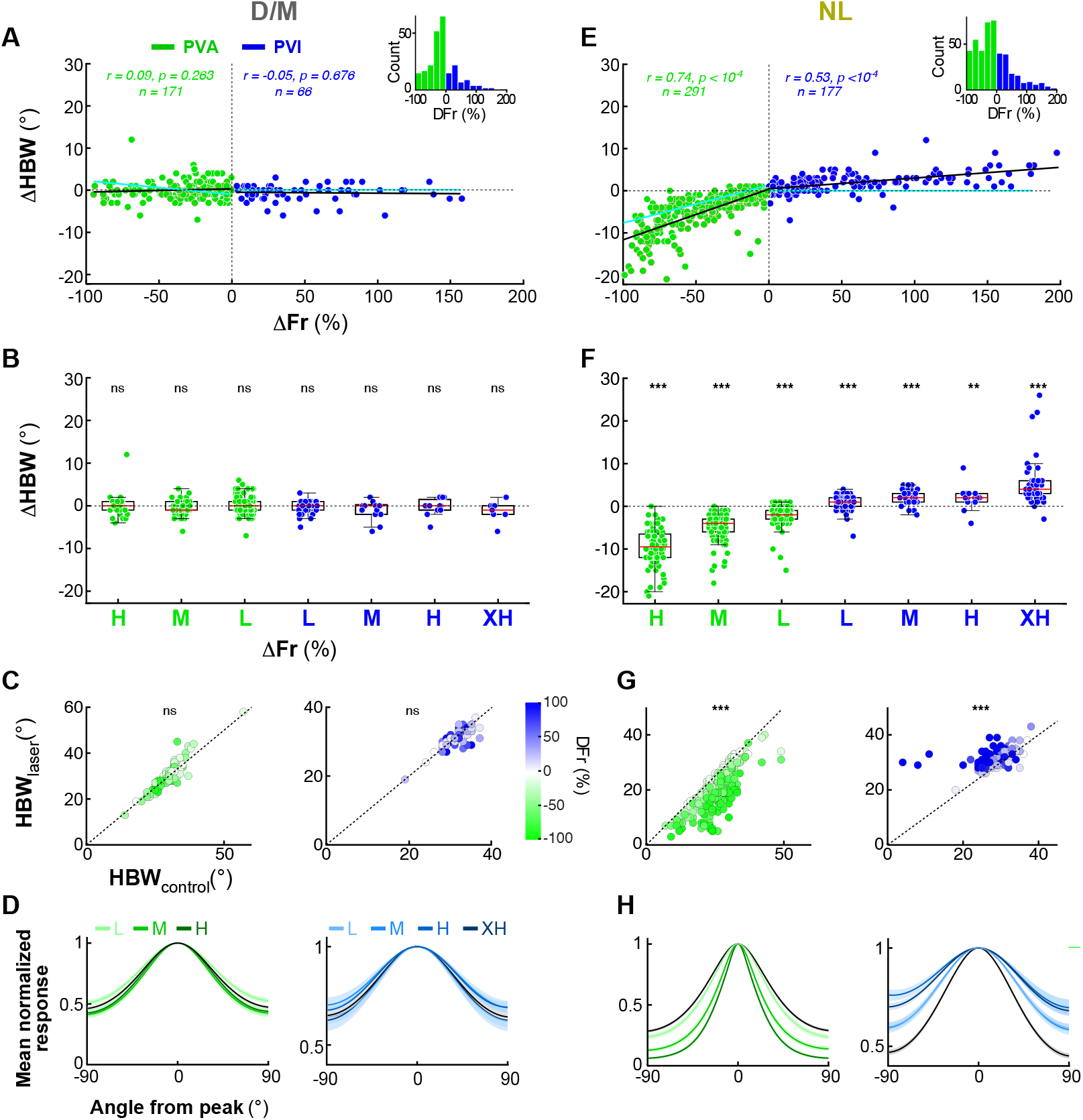
Distinct effects of *PV*^*+*^ interneuron manipulation on the orientation bandwidth of D/M and NL Pyr neurons’ tuning curves. **(A)** Scatter plot of the change in half-bandwidth (ΔHBW) as a function of the percent change in firing rate (ΔFr) for units classified as D/M. Here and in all remaining panels, *green dots* represent units recorded during PVA while *blue dots* indicate units in PVI; *dots outlined in red* are units with higher orientation selectivity (having circular variance, CV < 0.8). *Black solid line*: linear regression fit to the data (correlation, *r*, values and significance of the correlation, *p* values, are indicated at the top separately for PVA and PVI data). The number of cells (n) included in the analysis are also indicated at the top; note that here individual cells recorded at different intensity levels are counted as independent samples and presented as separate dots in each group, hence the n reported at the top is larger than the n in other analyses. *Cyan line*: linear regression fit of simulated tuning curves generated using the Threshold Linear Model^17^ (see STAR Methods, and **Fig. S6A, C**). *Inset*: Distribution of unit counts across the observed range of ΔFr. **(B)** The same data as in panel (A) is plotted as a box plot of ΔHBW for different percent changes in Fr: from low (L) to extra-high (XH) as described in the Results. *ns*: statistically non-significant changes in ΔHBW for that group (Wilcoxon signed-rank test). *Red lines*: medians. **(C)** Scatter plots of HBW in the laser condition vs. HBW in the control condition for PVA data (<u>Left</u>) and PVI data (<u>Right</u>). The relative ΔFr is represented as indicated by the color scale. **(D)** Average normalized population tuning curves in the control condition (*black*) and for each ΔFr group (same groups as in (B), from L to XH) indicated as different shades of *green* (for PVA data: <u>Left</u>) and different shades of *blue* (for PVI data; <u>Right</u>). Tuning curves were normalized to the peak responses in each condition. **(E–H)** Same as in panels (A–D), but for the units classified as NL. *Asterisks* in (F,G) indicate statistical significance according to the following convention: *= *p*<0.05, **=*p*<0.001, ***: *p*<0.0001. This convention also applies to other figures. Similar data for the Unclassified units is show in **Fig. S6F**.

The D/M Pyr cell population showed no significant change in HBW across all tested irradiance levels (**Fig. 2A**, *black regression line*, PVA: *r* =0.09, *p=* 0.2631, n= 171; PVI: *r* =-0.05, *p=* 0.6760, n= 66; Pearson correlation-for the purpose of this analysis the same cells recorded at different laser intensities are considered independent samples, hence the larger n). In **Fig. 2B** the same data are shown as box plots with cells grouped by magnitude of laser effect (ΔFr): Low (L: 0 to ±33% ΔFr), Medium (M: ±33 to ±66% ΔFr), High (H: ±66 to ±100% ΔFr), Extra-high (XH: +100 to ≥200% ΔFr). Again, there was no significant change in HBW within any magnitude group (p>0.11 for all within group comparisons; Wilcoxon signed-rank test). Scatter plots of HBW in laser vs control conditions also revealed no significant difference in HBW between these two conditions, for the D/M population (mean HBW±s.e.m for control vs laser = 29.8°±0.32 and 29.9°±0.38 for PVA, *p=*0.38; and 31.5°±0.35 and 31.3°±0.33 for PVI, *p=0*.24; Wilcoxon signed-rank test; **Fig. 2C**), as also evident in the similarity of the normalized population-averaged tuning curves between control and laser conditions, across tested irradiances (**Fig. 2D**). In contrast, the NL population showed a significant correlation between ΔHBW and ΔFr (PVA: *r* = 0.74, *p* < 10^34^, *n* = 291; PVI: *r* = 0.53, *p* < 10^34^, *n* = 177) (**Fig. 2E**). At all tested irradiances (L,M, H, XH) PVA significantly reduced HBW (i.e. sharpened orientation tuning), with ΔHBW = −9.56° ± 4.3 (*p* < 10^34^, *n* = 80; Wilcoxon signed-rank test) for the highest irradiance group, while PVI significantly increased HBW (broadened orientation tuning), with ΔHBW = 5.24° ± 4.9 (*p* < 10^34^, n=57) for the highest irradiance group (**Fig. 2F**). The difference between the mean HBW in control (PVA: 26.1°±0.36; PVI: 28.9°±0.27) vs. laser (PVA: 21.2°±0.4; PVI: 31.4°±0.2) conditions (pooled across irradiance levels) across the population was statistically significant (*p*<10^−5^ for both PVA and PVI, Wilcoxon signed-rank test) (**Fig. 2G**). The normalized population tuning curves showed a clear tuning sharpening for PVA, and broadening for PVI, which increased with the laser-induced magnitude of the ΔFr (**Fig. 2H**). The threshold linear model^17^, captured the effects of *PV*^*+*^ cell manipulations for the D/M population, but not for the NL population (**Fig. 2A,E**, *cyan fit*). This model predicts no change in HBW with PVI, but small decreases with PVA of higher intensity due to the “iceberg effect” (**Fig. 2E** *cyan fit*), the phenomenon by which Pyr neurons fail to reach spike threshold at orthogonal orientations due to a floor effect. In contrast to this model’s predictions, the NL population showed significant increases in HBW for PVI, and larger decreases for PVA than predicted by the iceberg effect, including at the lowest irradiance levels (**Fig. 2E**, compare *black fit with cyan fit*). The threshold linear model is described in detail in a subsequent section of the Results and in **Fig. S6**, which also shows the changes in HBW as a function of changes in firing rate for the Unclassified cell population).

There are different ways to measure orientation tuning and selectivity, each revealing different features of the orientation tuning curve^37^. HBW is a local measure of tuning sharpness around the peak of the tuning curve. Other useful measures are the orientation selectivity index (OSI) and the circular variance (CV) (see STAR Methods for how these were computed). The OSI is a global measure of orientation selectivity which compares responses at two widely separated values of orientation angle (the preferred and non-preferred orientations). OSI ranges between 0 (no selectivity) and 1 (highly selective). CV is another global measure of orientation selectivity that takes into account the shape of the tuning curve at all orientations; CV varies between 0 (highly selective curve) and 1 (flat curve).

We measured changes in OSI (ΔOSI) and CV (ΔCV) as a function of laser-induced percent changes in firing rate (ΔFr) separately for D/M and NL Pyr cell populations (**Fig. 3**). The D/M population exhibited non-significant or minimal changes in both OSI (**Fig. 3A-B**) and CV (**Fig. 3E-F**) across all ΔFr magnitude groups (L to XH), i.e. irrespective of laser intensity. Although statistically significant regressions were observed for OSI in both PVA and PVI conditions (**Fig. 3A**), and for CV in the PVA condition (**Fig. 3E**), effect sizes were modest. For example, for the largest effect magnitude group, ΔOSI = −0.01 ± 0.04 (*p* < 0.05, *n* = 103) for PVA, and ΔOSI = −0.04 ± 0.05 (*p* < 0.01, *n* = 11) for PVI; ΔCV = −0.01 ± 0.02 (*p* < 0.01, *n* = 42) for PVA, and ΔCV = 0.01 ± 0.01 (*p* < 0.05, *n* = 11) for PVI (**Fig. 3B,F**). In contrast, the NL population exhibited significant changes in both OSI and CV across all laser intensity levels for both PVA and PVI conditions, and these changes were significantly correlated with laser-induced changes in firing rates (ΔFr). Significant regressions (*black dashed lines* in **Fig. 3 C,G**) confirmed the systematic effect of *PV*^+^ manipulations on OSI and CV in this population (OSI–PVA: *r* = −0.70, *p* < 10^34^, *n* = 291; OSI–PVI: *r* = −0.76, *p* < 10^34^, *n* = 177; CV– PVA: *r* = 0.75, *p* < 10^34^, *n* = 291; CV–PVI: *r* = 0.74, *p* < 10^34^, *n* = 177). For the highest intensity group, PVA significantly increased OSI (ΔOSI = 0.32 ± 0.16, *p* < 10^34^, *n* = 80) and reduced CV (ΔCV = −0.34 ± 0.16, *p* < 10^34^; *n* = 80). Conversely, PVI significantly reduced OSI (ΔOSI = −0.31 ± 0.13, *p* < 10^34^, *n* = 57) and increased CV (ΔCV = 0.17 ± 0.14, *p* < 10^34^, *n* = 57) (**Fig. 3D,H**). For both the D/M and NL groups, the threshold linear model provided predictions of ΔOSI and ΔCV as a function of ΔFr that were consistent with the observed data (*cyan dashed lines* in **Fig. 3A,C,E,G)**. While OSI and CV capture the overall sharpening or broadening of the tuning curve, they are relatively insensitive to the detailed nonlinear “Mexican-hat” structure that distinguishes the NL from the D/M Pyr cell population, which instead significantly affects HBW. The threshold linear model fits to these data are discussed in detail in a subsequent section of the Results and in **Fig. S7**, which also shows the changes in OSI and CV as a function of changes in firing rate for the Unclassified cell population.

**Figure 3.**
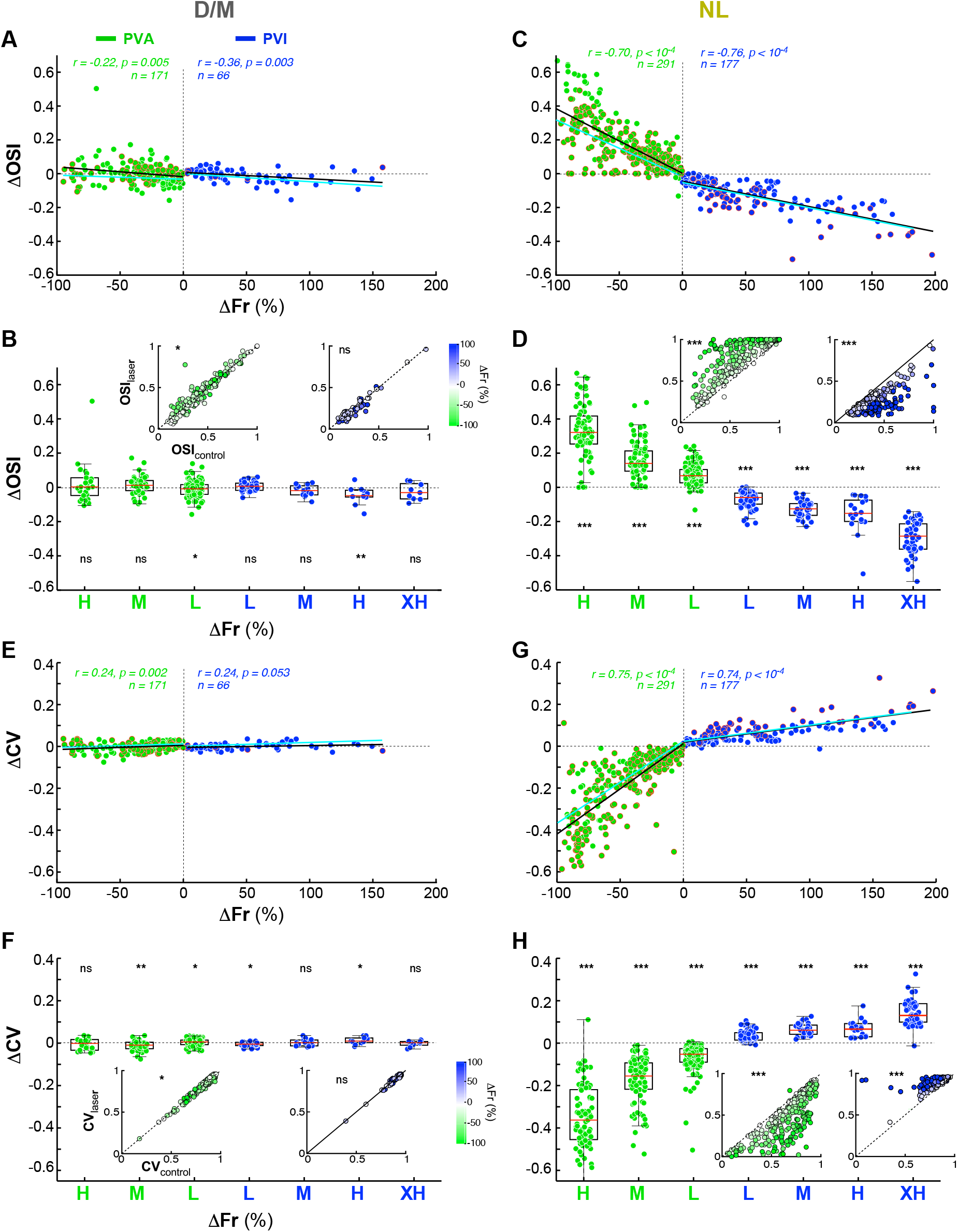
Distinct effects of *PV*^*+*^ interneuron manipulation on the orientation selectivity and circular variance of D/M and NL Pyr neurons’ tuning curves. **(A)** Scatter plot of the change in orientation selectivity index (*ΔOSI*) as a function of the percent change in firing rate (ΔFr) for units classified as D/M. Other conventions are as in **Fig. 2A**. The *cyan line* shows the linear regression fit of simulated tuning curves generated using the Threshold Linear Model (see STAR Methods and **Figs. S7A**). **(B)** Box plot of the same data as in panel (A) with conventions as in **Fig. 2B**. *Insets*: Scatter plots of OSI in the laser condition vs. OSI in the control condition for PVA data (<u>Left</u>) and PVI data (<u>Right</u>). The relative ΔFr is represented as indicated by the color scale. **(C,D)** Same analyses as in panels (A,B) but for units classified as NL. The *cyan line* shows the linear regression fit of simulated tuning curves generated using the Threshold Linear Model (see STAR Methods and **Fig. S7C**). **(E)** Scatter plot of the change in circular variance (*ΔCV*) as a function of the percent change in firing rate (ΔFr) for units classified as D/M. Other conventions are as in **Figs. 2A,3A**, but the *cyan line* refers to the simulated data shown in **Fig. S7G. (F)** Box plot of the same data as in panel (E) with conventions as in **Figs. 2B,3B**. *Insets*: Scatter plots of CV in the laser condition vs. CV in the control condition for PVA data (<u>Left</u>) and PVI data (<u>Right</u>). **(G,H)** Same analyses as in panels (E,F) but for units classified as NL. Here the *cyan line* refers to the simulated data shown in **Fig. S7I**. Similar data for the Unclassified units is show in **Fig. S7E,F,K,L**.

### Laminar-Specific Effects of *PV*^+^ Interneuron Manipulation on Pyr Neuron Response Gain and Orientation Tuning

Given the different effects of *PV*^*+*^ neuron manipulation on the D/M and NL Pyr neuron populations, we asked whether these two neuronal populations show any distinctive features, e.g. in their RF properties or laminar location. We used the laminar profile of multi-and single-unit spiking activity (**Fig. S4A-C**), current source density (CSD) (**Fig. S4D**), and local coherence spectrum (**Fig. S4E**), to assign single units to one of three laminar groups: supragranular (SG), G, or infragranular (IG) (see STAR Methods).

We found that D/M and NL units differed in RF properties: NL units exhibited significantly sharper orientation tuning (higher OSI, lower CV, and narrower HBW) than D/M units (**Fig. 4A-B**). For the NL units, the mean OSI (0.62±0.01 for PVA, 0.42±0.01 for PVI), mean CV (0.67±0.01 for PVA, 0.8±0.01 for PVI), and mean HBW (26.1°±0.36 for PVA, 28.9°±0.27 for PVI) differed significantly from those of D/M units (OSI: 0.41±0.01 for PVA, 0.25±0.02 for PVI; CV: 0.81±0.01 for PVA, 0.89±0.01 for PVI; HBW: 29.8°±0.30 for PVA, 31.5°±0.35 for PVI; *p<* 10^−9^ for all comparisons in **Fig. 4A,B**; Wilcoxon rank-sum test). Furthermore, as predicted, the D/M units were more prevalent in the G layer, whereas the NL units were more prevalent in the extra-G layers (**Fig. 4C**). Specifically, across the population of D/M and NL units, ~70% of units in SG and IG layers were NL, while 70% of units in G layer were D/M. **Fig. S5** shows the RF properties and laminar distribution for the Unclassified cell group, in comparison with those of the DM and NL groups.

**Figure 4.**
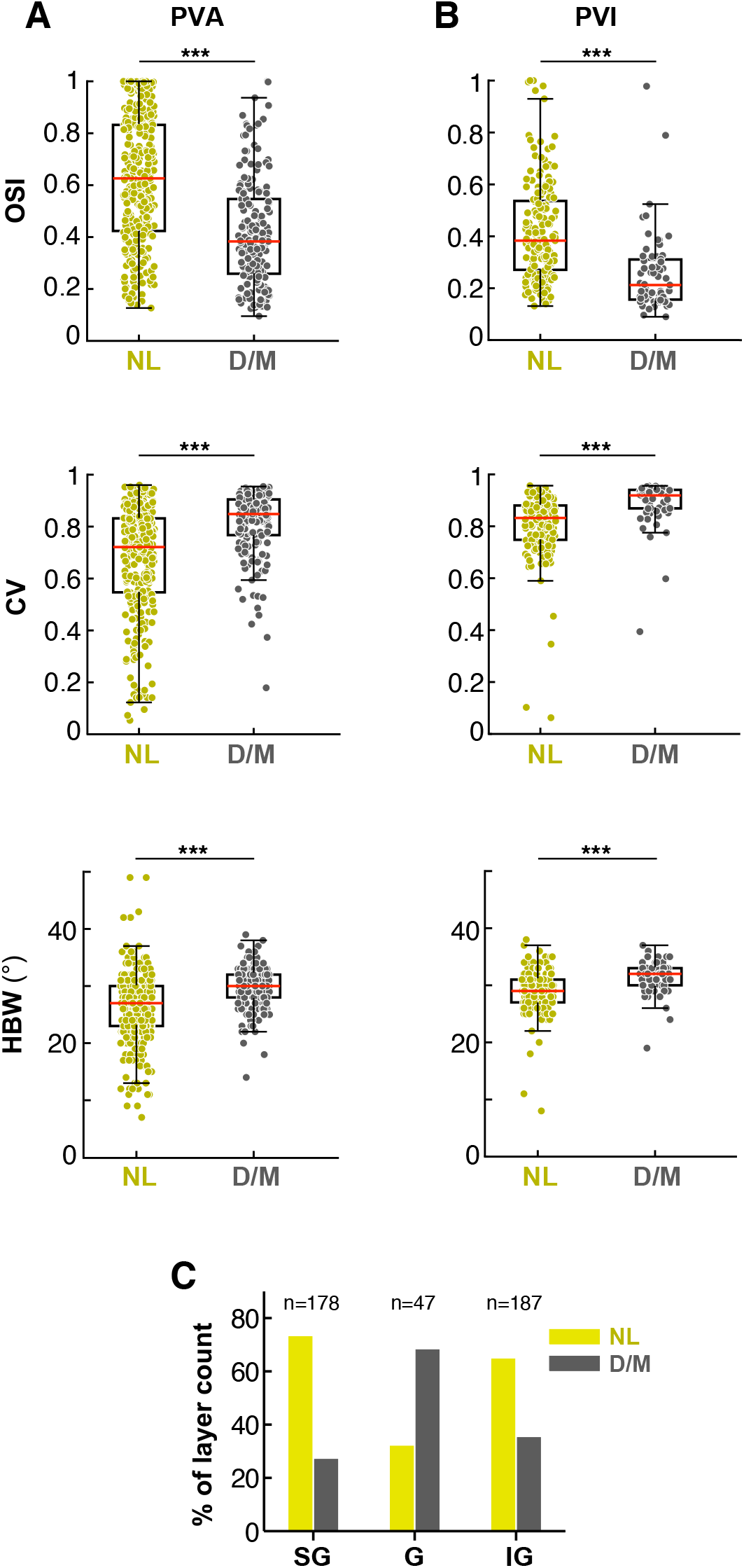
D/M and NL Pyr neurons have different tuning properties and laminar distribution. **(A)** Box plots of OSI (<u>Top</u>), CV (<u>Middle</u>) and HBW (<u>Bottom</u>) for NL (*yellow dots*) and D/M (*gray dots*) Pyr neurons recorded in the control (no laser) condition in PVA experiments. *Red horizontal lines*: median. **(B)** Same as in (A) but for units recorded in the control condition in PVI experiments. **(C)** Percent distribution of D/M vs NL units in different layers. n= total number of D/M + NL units in each layer (excludes U units). **Fig. S5** additionally shows the tuning properties and laminar distribution of Unclassified cells.

We next examined the impact of *PV*^+^ neuron manipulation on the orientation tuning curves of Pyr neurons residing in different V1 layers. This analysis was blind with respect to effect type (D/M, NL or U), therefore, it was performed on all orientation-selective units in our sample (n=495;see STAR Methods). **Figure 5A** shows the population average tuning curves of neurons in different V1 layers in the control (no laser) condition. In marmoset V1, as in macaque V1^25^, orientation tuning in the G layer is broader than in extra-G layers. As expected based on the prevalence of NL cells in extra-G layers, in both SG and IG layers, PVA reduced firing rates, increased orientation selectivity and sharpened orientation tuning (**Fig. 5B**), as evidenced by a significant increase in mean OSI [SG: mean ΔOSI± s.d. (pooled across laser intensities) = 0.092 ± 0.16, *p* < 10^34^, *n* = 200; IG: ΔOSI = 0.118 ± 0.16, *p* <, *n* = 327; Wilcoxon signed − rank test] and a reduction in mean CV and HBW (SG: ΔCV = −0.112 ± 0.16, *p* < 10^34^; ΔHBW = −3.07° ± 4.7, *p* < 10^34^, *n* = 200; IG: ΔCV = −0.11 ± 0.15, *p* < 10^34^; ΔHBW = −2.76° ± 4.34, *p* < 10^34^, *n* = 327) (**Fig. 5C-E**; for the purpose of these analyses the same cells recorded at different intensities are counted as independent samples). In contrast, PVA reduced firing rate, but did not induce a significant change in orientation selectivity and tuning in the G layer (**Fig. 5B-E**). Conversely, PVI increased firing rates and broadened the orientation tuning curve in SG and IG layers (**Fig. 5F**), as also indicated by a significant decrease in mean OSI and increases in mean CV and mean HBW in these layers (SG: n= 132; ΔOSI = −0.14 ± 0.12, *p* < 10^34^; ΔCV = 0.069 ± 0.06, *p* < 10^34^; ΔHBW = 1.76° ± 2.33, *p* < 10^34^. IG: *n* = 100; ΔOSI = −0.12 ± 0.15, *p* < 10^34^; ΔCV = 0.06 ± 0.13, *p* < 10^34^; ΔHBW = 1.7° ± 4.2, *p* < 10^34^; Wilcoxon signed-rank test) (**Fig. 5G–I**). However, PVI did not significantly affect tuning sharpness (HBW) in the G layer (**Fig. 5I**). Although statistically significant changes in OSI and CV were observed in the G layer, the magnitude of these changes was small (ΔOSI = −0.06 ± 0.09, *p* < 0.005; ΔCV = 0.02 ± 0.04, *p* < 0.05; *n* = 33; **Fig. 5G − H**). Notably, PVI made the tuning HBW of extra-G layer neurons resemble that of G layer cells in the control condition (**Fig. 5I**). This result supports the hypothesis that *PV*^*+*^ neuron-mediated inhibition sharpens orientation tuning outside the G layer. In contrast, the observation that *PV*^*+*^ cell manipulations in the G layer affect Pyr neuron firing rates (**Fig. 5B,F**), without altering orientation tuning width supports a role for *PV*^*+*^ interneurons in this layer in gain control. These results are consistent with the observation that cells showing D/M effects of *PV*^*+*^ cell manipulation dominate in G layers whereas those showing NL effects are most numerous in the extra-G layer (**Fig. 4C**).

**Figure 5.**
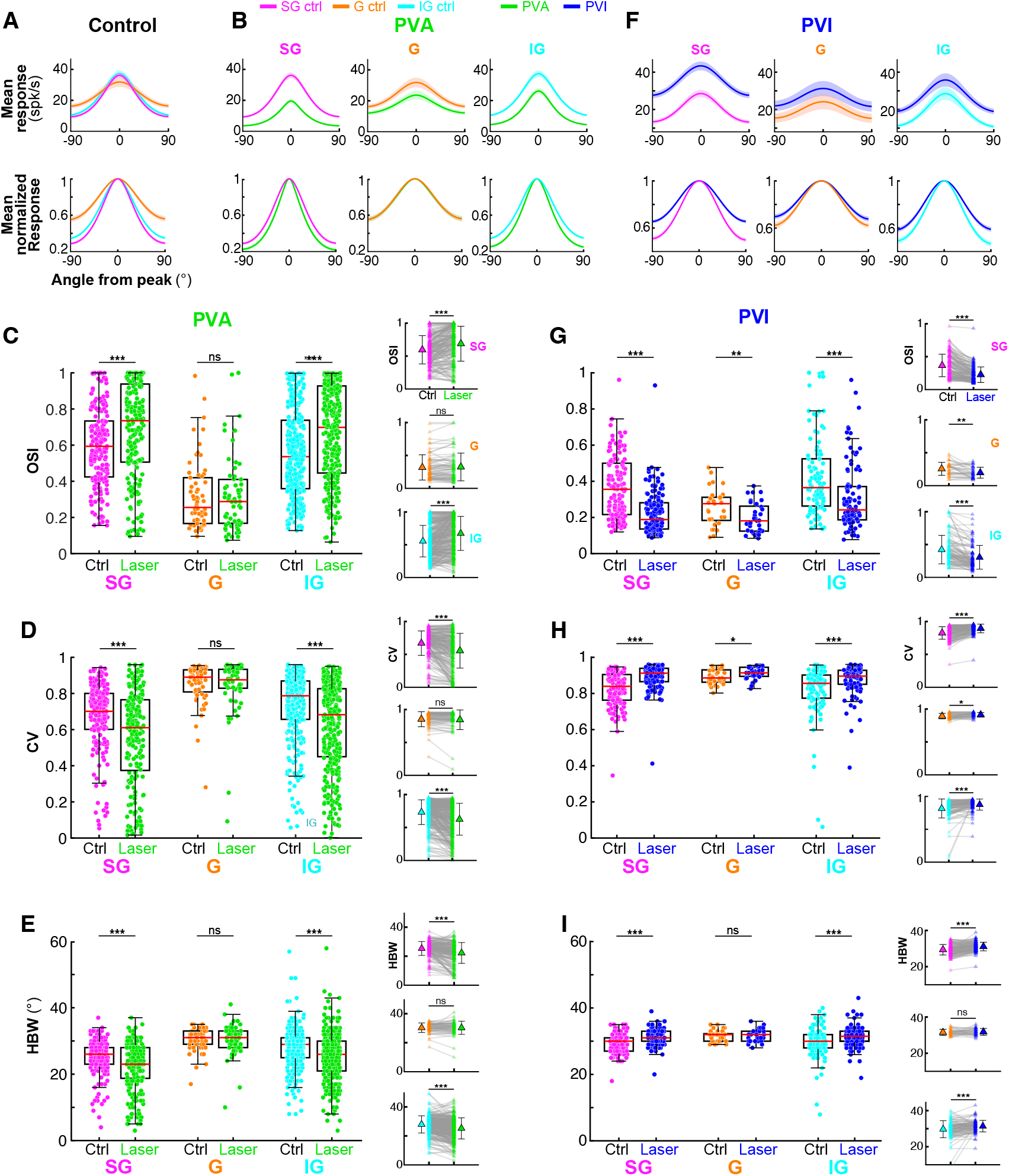
Laminar-specific effects of *PV*^*+*^ interneuron manipulation on Pyr neuron response gain and orientation tuning. **(A)** Average population tuning curves in the control condition for SG (*pink*), G (*orange*) and IG (*cyan*) units. Here and in panels (B,F), <u>Top</u>: mean firing rate, and <u>Bottom:</u> mean firing rate normalized to the peak in each layer. Shading: s.e.m. **(B)** Average population tuning curves in control (*pink, orange, cyan*) and PVA (*green*) conditions for each layer group. **(C-E)** <u>Left</u>: Box plots of the distributions of OSI (C), CV (D), and HBW (E) in control and PVA conditions for each layer group. <u>Right:</u> pairwise comparisons (control vs. laser) for each unit in the different layer groups. **(F-I)** Same as (B-E) but for PVI (*blue*).

### Theoretical Models

Previous mouse studies of the effect of *PV*^*+*^ neuron manipulation on Pyr neuron responses reported apparently conflicting results. In one study, *PV*^*+*^ neuron activation changed Pyr neurons’ firing rates divisively, affecting response gain but not tuning^16^. Another study reported that PVA subtractively lowers Pyr neuron responses, sharpening orientation tuning^15.^ A third study reported mixed divisive/multiplicative and subtractive/additive effects, which affected response gain but only minimally tuning^17^. However, it was argued that all effects observed in these previous studies could be explained by a threshold linear model (TLM) with both multiplicative/divisive and additive/subtractive components^36^; this model predicts small tuning changes when *PV*^*+*^ neurons are photoactivated at high intensities, as observed by^15^, due to an “iceberg effect”^14^ but no changes in tuning when *PV*^*+*^ neurons are photoinactivated at any intensity.

To evaluate whether the TLM can account for our observed D/M and NL effects of *PV*^*+*^ neuron manipulation, we simulated the effects of laser manipulations on neuronal orientation tuning curves based on this model (see STAR Methods), and then analyzed changes in HBW, OSI and CV as a function of ΔFr for the simulated data (**Fig. S6–S7**), as done in **Figs. 2-3** for the real data. For the D/M population, the TLM model provided good fits to the neuronal orientation-tuning curves under the laser condition (goodness-of-fit, median R^2^= 0.87±0.01), and accurately predicted changes in HBW, OSI and CV (**Figs. S6A-B, S7A,B,G,H**). In contrast, for the NL population, the TLM, although providing reasonably good fits to the neuronal orientation-tuning curves under laser conditions (median R^2^= 0.92±0.006), failed to accurately predict changes in HBW, especially those induced by PVI, while also underestimating the effects of PVA (**Fig. 6SC-D**). Notably, in the data, the PVA-induced sharpening of tuning curves was evident even at laser intensities that produced minimal reductions in firing rate (**Figs. 2E,F, S6D**). In contrast, in the TLM-simulated data, sharpening of tuning curves only occurred at laser intensities causing ≥ 30% reduction in firing rate, and the magnitude of sharpening was smaller than observed in the data (**Fig. S6C,D**). This is because the TLM produces tuning sharpening via an iceberg effect, i.e. once the response at non-preferred orientations reaches zero. Changes in OSI and CV for the NL population, instead were better captured by the TLM (**Figs. S7C,D,I,J**). **Figs. S6E-F** and **S7E,F,K,L** show TLM-simulations and real data for the Unclassified cell population.

**Figure 6.**
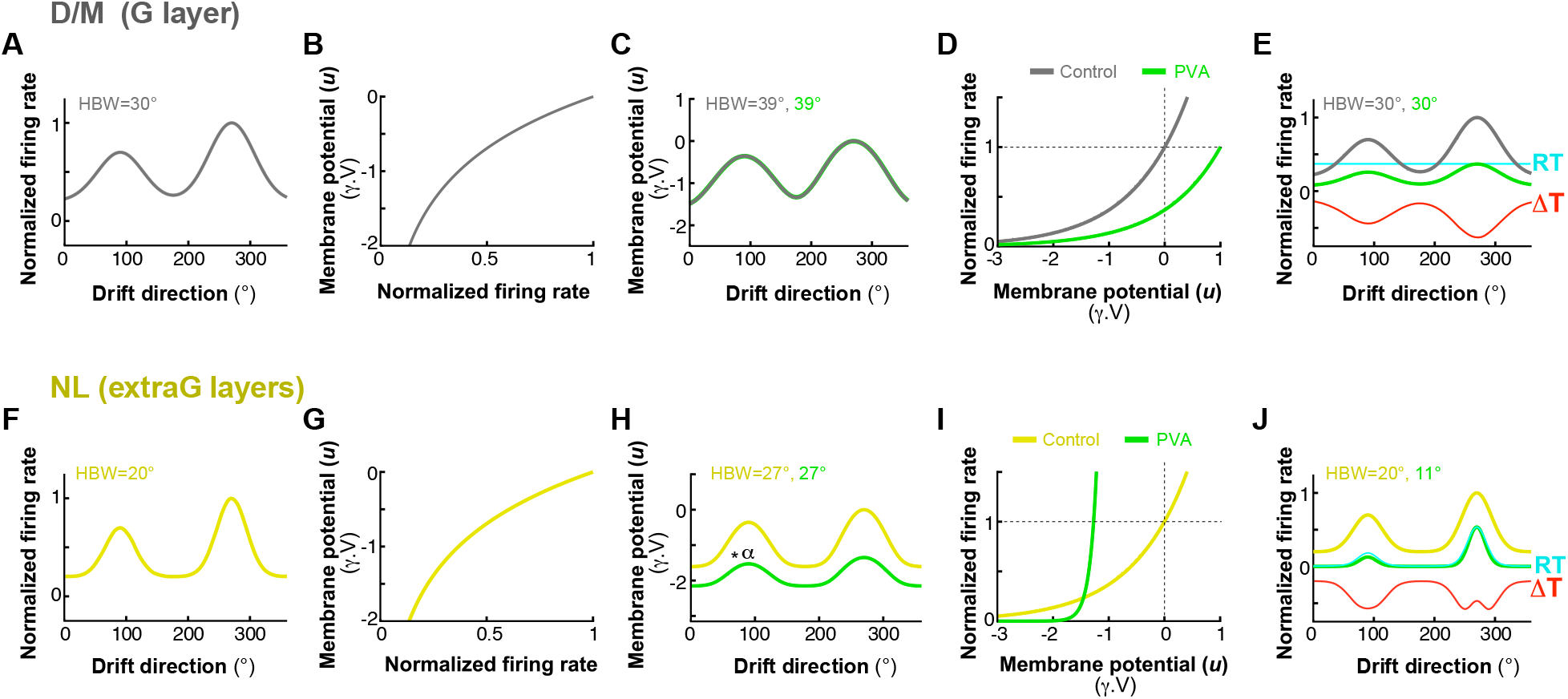
The I/O model. The I/O model for D/M cells in the G layer (A-E) and for NL cells in the extra-G layers (F-J). **(A)** Example normalized firing rate tuning curve of a D/M cell from the data, representing the model’s starting point. **(B)** Reversed I/O function, where the x-axis is the normalized firing rate and the y-axis is membrane potential in arbitrary voltage units (*γV*), determined by the chosen *ϑ* and *β* values. **(C)** Membrane potential tuning curve in the control condition (*gray curve*), *T*_*MC*_, computed as *T*_*M*C_ = *f*^−1^(*T*_Fr*C*_), where *T*_*FrC*_ is the firing rate (output) tuning curve in the control condition. Here the laser-induced membrane potential tuning curve (*green curve*), *T*_*ML*_, is considered identical to *T*_*MC*_, as the driving LGN input is not affected by the intracortical *PV*^*+*^ neuron manipulations. **(D)** I/O function used to transform the membrane potential tuning curve, *T*_*M*,_ into the corresponding firing rate (output) tuning curve, *T*_*Fr*_. The *gray curve* shows the control I/O function with the same parameters as used in the reversed I/O function (B); the *green curve* represents a rightward-shifted I/O function (due to increased *ϑ*), caused by the laser manipulation (in this example, PVA). (**E)** Model firing rate output for each I/O function. The *gray* tuning curve is identical to the one in (A) (generated with identical I/O parameters). The *green* tuning curve results from the shifted I/O function (*green* in D), reproducing the multiplicative effect observed in the data for the D/M population. As in **Fig. 1B,D**, the *red curve* (*ΔT*) is the difference between the green and gray curves, and the *cyan line* (*RT*) *is* the ratio between these same curves, which is a constant value (multiplicative gain). **(F,G**) Same as (A,B) but the example tuning curve in (A) is of a NL cell. **(H)** The *yellow curve* is the membrane potential tuning curve in the control condition, *T*_*MC*_, computed as in panel (C). The *green curve* is the membrane potential tuning curve in the laser condition, *T*_*ML*_, which is a multiplicatively-scaled version (*α* ratio) of *T*_*MC*_, as the *PV*^*+*^ neuron manipulation causes a D/M effect on the driving G layer input to neurons in the extra-G layers. **(I)** I/O function. *Yellow curve*: control I/O function with the same parameters as used in the reversed I/O function (G). Increasing *β* and reducing *ϑ* in the I/O function causes a leftward shift and increases the steepness of the function (*green curve*), reproducing the NL effects of PVA seen in the firing rate tuning curve in our NL population **(J)**.

Given the failure of the TLM to fully account for the observed NL effects, we developed a model to better capture the broad range of *PV*^+^ manipulation effects on orientation tuning and response gain (**Fig. 6**). Neuronal responses can be modeled as input/output (I/O) functions (graphed in e.g. **Fig. 6D**), which describe how the membrane potential (cell’s input) is transformed into the spike rate (cell’s output). This relationship is non-linear and well captured by a simple exponential function, which can take several forms^14,38-40^, and is modulated by factors such as stimulus condition, and input noise^40-46^. We adopted the “escape-rate” formulation of this function, which has a strong theoretical foundation in neural dynamics^40,47,48^:

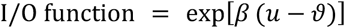

where *u* is the cell’s membrane potential, *ϑ* is the cell’s firing rate threshold, and *β* controls the rate of exponential growth (how rapidly the probability of firing rises as *u* approaches *ϑ*). In this model the neuron can fire even if the voltage is below the threshold, due to noise. The parameter *β* effectively controls the steepness of the neuron’s firing probability. For higher *β* values (steeper curve, low noise), the neuron’s firing probability increases for small changes in the membrane potential near or above threshold, but it decreases when the membrane potential is below threshold. Essentially, for higher *β* values, the neuron becomes almost deterministic, i.e. it fires almost exactly when *u* reaches threshold (“hard” threshold). For lower *β* values (shallower curve, higher noise), the firing probability increases gradually (“soft” threshold), the neuron can fire even when *u* is well below threshold, due to noise fluctuations.

Given an orientation tuning curve recorded in the control condition, *T*_*FrC*_, (e.g. **Fig. 6A**), using the inverse of the I/O function (**Fig. 6B**, with *ϑ* = 0 and *β* = 1), we can derive the membrane-potential tuning curve, *T*_*MC*_ (**Fig. 6C**). The absolute values of *ϑ* and *β* do not alter the shape of the membrane-potential tuning curve, rather, they only shift it vertically or scale it by a constant factor, respectively. Therefore, while the exact values of *ϑ* and *β* can be chosen arbitrarily, what is meaningful is their relative change caused by *PV*^*+*^ neuron manipulation. For D/M cells, which are prominent in the G layer and receive direct LGN inputs, we assume that, since our optogenetic manipulations do not affect LGN inputs but only intracortical cells, the membrane-potential tuning curve (largely the results of driving LGN inputs) is unchanged in the control and laser condition (**Fig. 6C**). With this assumption, we show that increasing the threshold *ϑ* reproduces the D/M effect of *PV*^*+*^ neuron manipulations on the output (i.e. firing rate) orientation tuning functions seen in our D/M population (**Fig. 6E**). While intracortical inputs also affect the membrane potential, here we assume their impact is small relative to that of geniculate inputs. Importantly, however, this assumption does not fundamentally change the model results, as using a scaled version of the control membrane potential simply shifts the I/O function along the input (*u*) axis. Note that the effect of increasing *ϑ* on the I/O function is a subtractive (rightward) shift along the input (*u*) axis (**Fig. 6D**), because in the I/O function, increasing *ϑ* subtracts a larger value from the membrane potential *u*. However, while the shift in the input space is subtractive, mathematically, its impact in output space (the firing rate) is a constant multiplicative scaling, due the exponential transformation (**Fig. 6E**).

For N/L cells, which are prominent in the extra-G layers and receive driving G layer input, instead, the model includes an additional step. Namely, it multiplies the membrane-potential tuning curve in the control condition, *T*_*MC*_ (the yellow curve in **Fig. 6H**), by a gain factor (*α*) to obtain the membrane-potential tuning curve in the laser condition, *T*_*ML*_, (green curve in **Fig. 6H**). This assumption is based on our own results showing that the G layer cells, which send inputs to the extra-G layer cells, show D/M effects (gain changes) of *PV*^*+*^ neuron manipulation. Specifically:

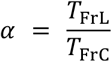

where, *T*_*FrL*_ and *T*_*FrC*_ are the firing rate tuning curves (output) in the laser and control condition, respectively. We obtain *α* from the empirical data by measuring laser-induced changes in firing rate (Δ*Fr*) in the G layer. We derive the resting membrane potential from the neuron’s spontaneous firing rate, using the inverse I/O function, subtract this from the *T*_*MC*_, and multiply the result membrane-potential tuning curve by *α*. Thus, under the assumption that the membrane potential for the cells in the extra-G layer is scaled divisively/multiplicatively by the *PV*^*+*^ neuron manipulation, we show that the combined effect of increasing *β* and reducing *ϑ* reproduces the NL effect of *PV*^*+*^ neuron manipulations on the output orientation tuning functions seen in our NL population (**Fig. 6J**). Theoretical and intracellular studies have demonstrated that this phenomenon (i.e. the increase in *β* value) is equivalent to reducing input noise^40,49,50^. Increasing *β* in the I/O function increases the steepness of the curve (i.e. the curve rises more sharply around the threshold *ϑ*) (**Fig. 6I**) and makes the neuron’s firing more deterministic and dependent on reaching a specific voltage threshold (due to lower noise). This leads to a lower average firing rate in the output tuning curve (**Fig. 6J**), because the neuron only fires when the membrane potential reliably crosses the threshold, which happens less frequently, thus lowering average firing rate. Moreover, this “suppressive” effect on firing rate is stronger for inputs evoking membrane potentials below the resting membrane potential (such as stimuli of non-preferred orientation), because the reduction in firing probability is strongest in this regime. In contrast, at the preferred orientation, the membrane potential is high, close to or exceeding threshold, so increasing *β* makes the response more reliable and potentially increasing firing rate even further. In other words, the escape function amplifies differences in subthreshold inputs when one input is close to threshold and the other is far away. As a result, the effect of increasing *β* is to sharpen the orientation tuning curve, by suppressing responses more strongly at the non-preferred orientations than at the preferred orientation.

### Fitting the I/O Model to the Data

We next sought to quantify the I/IO model’s ability to account for observed changes in orientation tuning. To this goal we fitted this model to the data.

We used *T*_highFr_ (the control tuning curve in PVA and the laser tuning curve in PVI) to compute the membrane potential tuning curve *(T*_*M*_). By fitting the I/O function that best transforms T_M_ into *T*_lowFr_ (the laser tuning curve in PVA and control curve in PVI), we extracted the corresponding I/O function parameters. The resulting *ϑ* and *β* values were then plotted as a function of the percent change in firing rate (**Fig. 7A-B**). The results indicate that for the D/M population, the magnitude of the *PV*^*+*^ neuron manipulation was positively (note here more negative values on the x axis correspond to larger decreases in firing rate, i.e. higher *PV*^*+*^ neuron manipulation magnitude) and significantly correlated with the value of *ϑ* (**Fig. 7A** *gray dots*, **Fig. 7C**), whereas the value of *β* did not change with the magnitude of the manipulation (**Fig. 7B** *gray dots*, **Fig. 7C**). Instead, for the NL population, the magnitude of the *PV*^*+*^ neuron manipulation was positively and significantly correlated with the value of *β* (**Fig. 7B** *yellow dots*, **Fig. 7D**) and negatively and significantly correlated with the value of *ϑ* (**Fig. 7A** *yellow dots*, **Fig. 7D**). This suggests that *PV*^*+*^ neuron manipulation can lead to D/M effects by changing the neuron’s spiking threshold, and to NL effects by changing both threshold and the steepness of the I/O curve (i.e. by changing input noise).

**Figure 7.**
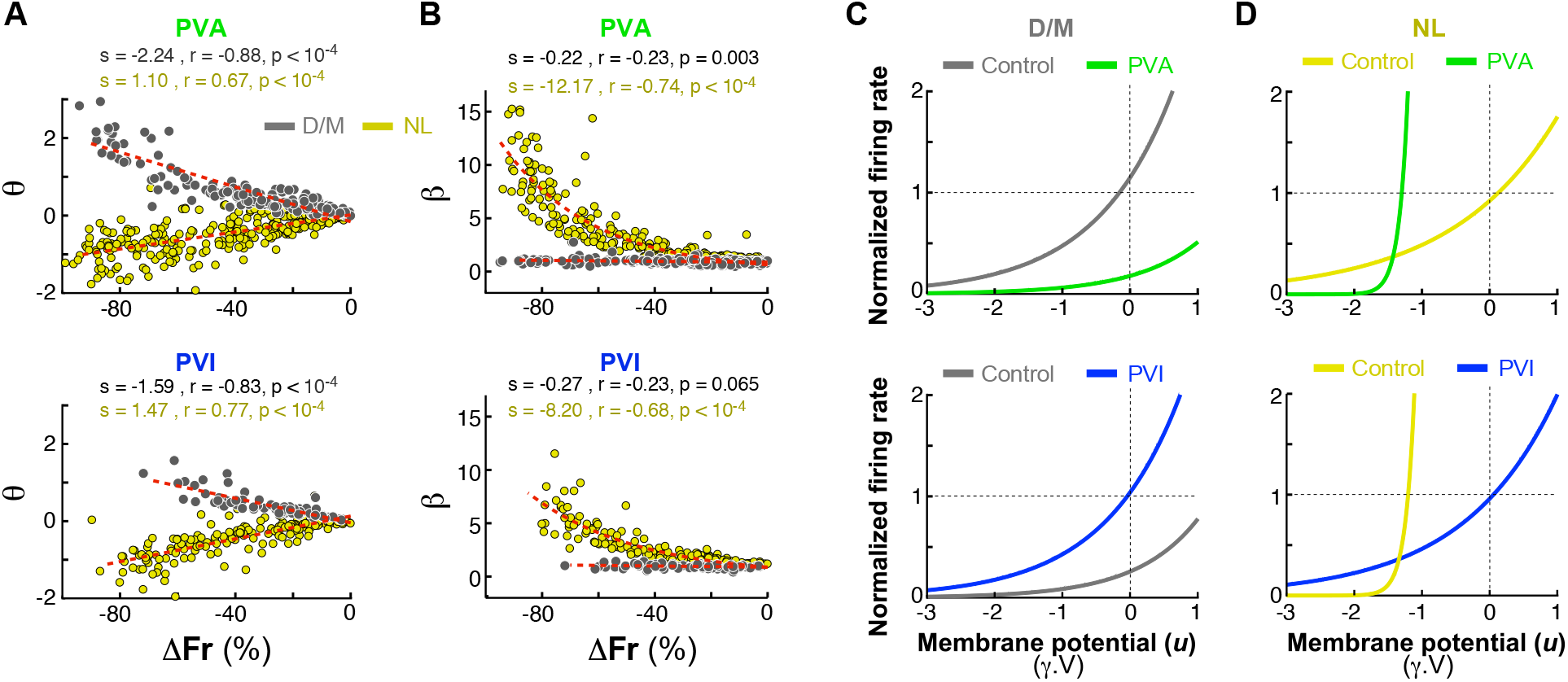
I/O model evaluation and parameter calculation. **(A,B)** Estimated I/O function parameters—threshold (*ϑ*) (A) and exponential growth rate (*β*) (B)—for PVA (<u>Top</u>) and PVI (<u>Bottom</u>), plotted against *ΔFr. Gray dots*: D/M population; *yellow dots*: NL population. In PVI experiments, the lower *PV*^+^ neuron activity condition (i.e. the laser condition) was equivalent to the control in the PVA condition (therefore, *ΔFr* values are more negative with increasing magnitude of the manipulation), ensuring that *ΔFr* values fall within the same range (0% to −100%) as in PVA experiments. **(C,D)** I/O functions in control and laser condition for PVA (<u>Top</u>) and PVI (<u>Bottom</u>), for the D/M (C) and NL (D) populations. *Gray and yellow curv*es correspond to the control conditions, and *green and blue curves* to the PVA and PVI condition, respectively, as in the experiments (but as pointed out above, for modeling purposes, in PVI the control condition was treated as the blue curve).

Using the I/O model’s fitted parameters and the control *T*_highFr_, we then simulated the laser-induced *T*_lowFr_. Using the control (T_highFr_) and simulated laser (*T*_lowFr_) tuning curves, we then analyzed changes in HBW, OSI and CV as a function of ΔFr for the I/O model-simulated data (**Fig. 8**), as done in **Figs. 2-3** for the real data, and in **Fig. S6-S7** for the TLM model-simulated data. The I/O model, which has an equivalent number of free parameters (2) as the TLM, provided good fits to the neuronal orientation-tuning curves under the laser condition (for the D/M population, median R^2^=0.87±0.01, no significant difference with the TLM, p=0.72 Wilcoxon signed-rank test; for the NL population, median R^2^=0.93±0.006, and this value was significantly higher than the TLM, p<10^−17^ Wilcoxon signed-rank test), and, unlike the TLM model, accurately predicted changes in HBW, as well as OSI and CV, for both the D/M and NL populations (**Fig. 8**), and for the Unclassified cell population (**Fig. S8**).

**Figure 8.**
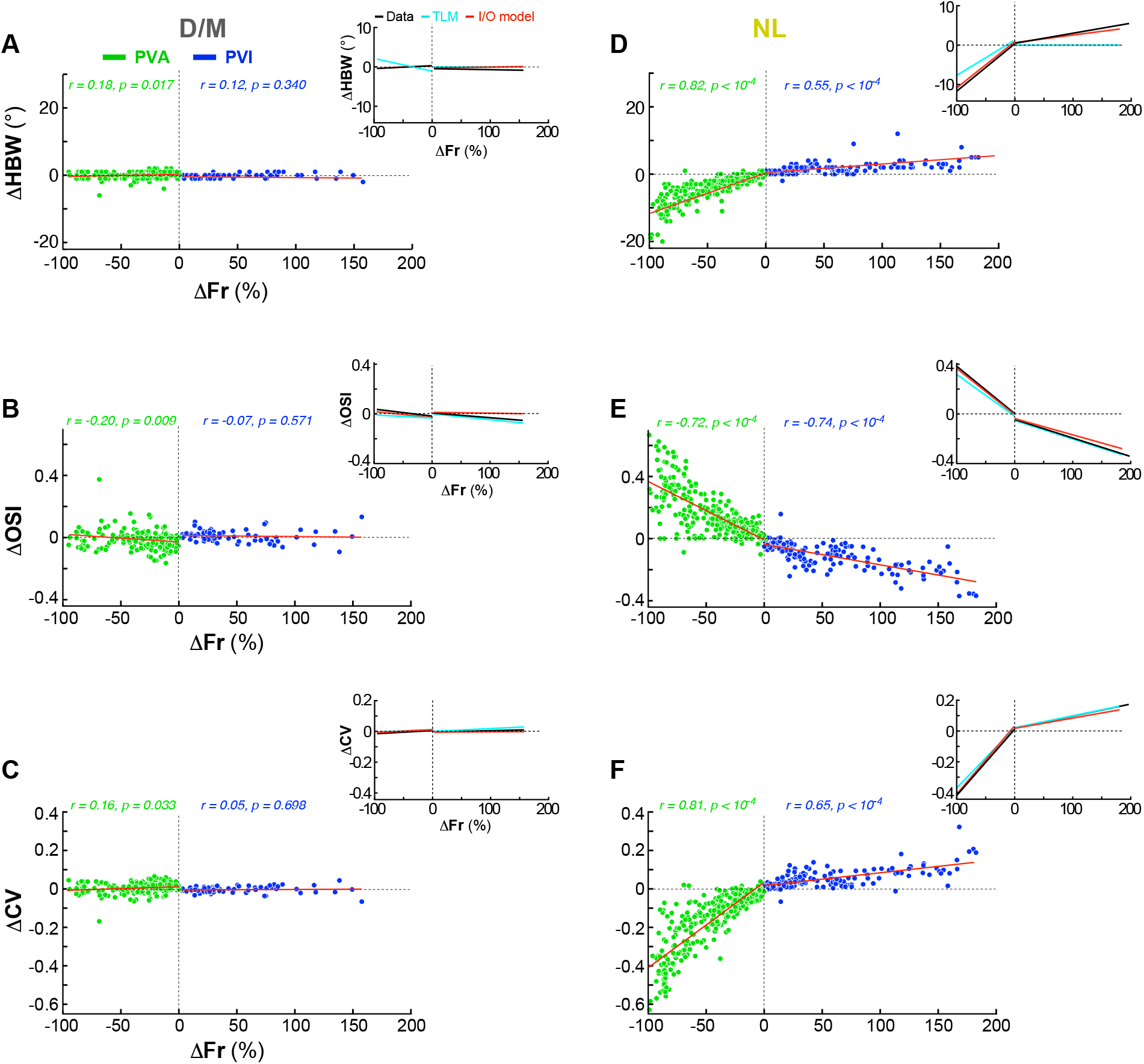
The I/O model captures the effects of PV^+^ neuron manipulations on orientation tuning and selectivity. **(A)** Scatter plot of the change in HBW as a function of the percent change in firing rate for I/O model-simulated effects of *PV*^*+*^ neuron manipulations on the D/M population. Here and in all remaining panels, the *red line* is the linear regression fit to the I/O-model-simulated data, to which the *r* and *p* values refer. *Inset*: comparison of linear regression fits to the data (*black line*), the TLM-simulated data (*cyan line*) and the I/O model-simulated data (*red line*). **(B)** Same as in (A) but for changes in OSI for the D/M population. **(C)** Same as in (A) but for changes in CV for the D/M population. **(D-F)** Same as (A-C) but for the NL population. Similar data for the Unclassified cell population is shown in **Fig. S8**. Other conventions are as in **Figs. 2-3**.

## Discussion

Using laminar array recordings and selective optogenetic manipulation of *PV*^*+*^ interneurons in marmoset V1, we found that the function of this inhibitory neuron class is layer-specific. In the geniculate input layer of V1, *PV*^*+*^ interneurons linearly control the response gain of pyramidal cells, but outside this layer, they non-linearly control pyramidal cells’ orientation tuning.

The role of inhibition in sharpening orientation tuning in carnivores and primates has been debated for decades. Two main theoretical solutions have been proposed for the emergence and refinement of orientation selectivity in area V1 of these species: feedforward models that rely on thalamocortical drive and intrinsic nonlinearities of cortical neurons (such as spike threshold, contrast saturation and spike rectification), and recurrent network models in which intracortical inhibition sharpens weakly orientation-biased geniculate inputs^12-14,37,51^. In the former models, inhibition primarily acts as a gain controller, while in the latter models it serves to sharpen sensory tuning. Here we show that one class of inhibitory neurons, the *PV*^*+*^ cells, can do both. In the granular layer, where thalamocortical afferents enter the cortex, and pyramidal neurons are weakly orientation-tuned, *PV*^*+*^ cells mainly act as gain controllers. As information flows from the granular to the extra-granular layers, pyramidal cells become more sharply orientation-tuned, and this tuning is mediated by *PV*^*+*^ cells in these layers.

Layer-specific *PV*^*+*^ neuron functions could result from distinct *PV*^*+*^ neuron subtypes and/or laminar-specific *PV*^*+*^ neuron connectivity. There is evidence for both hypotheses. Transcriptomic studies in primates have identified multiple subtypes of *PV*^*+*^ neurons^6,52^, often enriched in distinct layers. Moreover, *PV*^*+*^ cells comprise two main morphological subtypes, the basket and chandelier cells ^6,7,53^, having distinct laminar distribution. Even within the same morphological class, there exists layer-specific connectivity. For example, in primate V1, one type of basket cell, the wide-arbor basket cell, makes long-range trans-columnar axonal connections only outside the granular layer^54,55^. This cell type can provide spatially distributed, broadly-tuned inhibition to neurons in different orientation columns, preferring similar as well as different stimulus orientations. Recurrent models of orientation tuning rely on this type of broadly-tuned inhibition for sharpening weakly-biased tuning resulting from the selective convergence of non-orientation selective afferents from the thalamus or other cortical layers^13,37,56-58^. In contrast, basket cells in the granular layer make short local axonal connections with neurons within the same orientation column^59^, and may be better suited to mediate gain control. Gain control, a linear scaling of responses at all orientations, requires stronger *PV*^*+*^ inhibition at the preferred orientation of the suppressed cell, because excitatory conductance is higher at the preferred orientation. This could be implemented by broadly-tuned *PV*^*+*^ cells maximally suppressing local neurons having similar preferred orientation. Intra-columnar PV cells in the granular layer seem well positioned to carry out this task in the layer where feedforward thalamic afferents first enter the cortex. In support of these mechanisms is also evidence in mouse V1 that *PV*^+^ cells are untuned or more broadly orientation-tuned than excitatory neurons^16,17,60,61^. Recent evidence in primate V1 also shows that, despite being tuned for visual stimulus features, *PV*^*+*^ cells are more broadly tuned than excitatory neurons within the same orientation column^62^.

To explain the mechanisms underlying the different effects of *PV*^+^ cells in different layers, we investigated how inhibition could differentially affect the input/output (I/O) function of V1 cells^39,40^. We showed that a change in spike threshold (the parameter *ϑ* in the I/O model) can lead to a subtractive/additive shift of the I/O curve along the x-axis, producing a divisive/multiplicative effect of the output firing rate tuning curve (gain control). How could *PV*^*+*^ cells change a neuron spike threshold? An increase (decrease) in inhibitory conductance caused by *PV*^*+*^ neuron activation (inactivation), shifts the membrane potential further from (closer to) the threshold, so that a stronger (weaker) input would be required to reach threshold. Ultimately the neuron behaves as if the threshold is higher. Consistent with this hypothesized mechanism, intracellular recordings in mouse V1 have demonstrated that optogenetic inactivation of ^+^*PV* cells causes a reduction in direct synaptic conductance^17^, ultimately behaving as a reduction in threshold.

With respect to the non-linear effects of *PV*^+^ cell manipulation, which dominate in the extra-granular layers, we showed that an increase in the steepness of the I/O function (the parameter β in the I/O model), equivalent to reducing input noise^40^, coupled with a small decrease in spike threshold *ϑ*, makes the neuron’s firing more deterministic and dependent on reaching a specific voltage threshold. This ultimately leads to a reduction in the output firing rate, which is stronger at the non-preferred than preferred orientations, thus sharpening the orientation tuning curve. Mechanistically, changes in *PV*^*+*^ interneuron activity could alter noise levels in several ways. For example, inhibition can reduce trial-to-trial variability and shared (correlated) noise^63,64^, e.g. by reducing recurrent excitatory feedback amplifications and stabilizing cortical network dynamics^65^. Fast *PV*^+^-mediated inhibition also leads to quicker decay of noise fluctuations, shortening noise correlation times, with consequent reduced likelihood of threshold crossing^66^. *PV*^*+*^ interneurons targeting the perisomatic compartments of pyramidal cells, are well-suited to control input noise.

Synaptic noise and trial-to-trial variability in membrane potential increase under weak-signal conditions (e.g., at low stimulus contrast). Because trial-to-trial variability is partly responsible to carry the membrane potential above threshold, increased variability leads to increased spiking even when mean membrane potential is unchanged^41^, enabling reliable transmission of weak inputs and maintaining information^40^. Instead at high contrast, noise and variability decrease, leading to a decrease in spiking for the same mean membrane potential, sharpening of feature tuning, and improved fidelity of transmitted information. This contrast-dependence of trial-to-trial variability of the V1 neurons’ membrane potential, which results from the response variability of LGN afferents^67^, has been proposed to underlie the contrast-invariance of orientation tuning in cat V1 cells^41^. Our results indicate that in primate V1 intracortical inhibition is indispensable for sharpening the tuning and selectivity of pyramidal cells, and suggest that the strength of this inhibition may be dynamically controlled by feedforward inputs, thereby supporting and integrating elements of both feedforward and recurrent models.

Our results reconcile conflicting results from previous rodent studies on the impact of *PV*^+^ neuron manipulation on the orientation tuning function of pyramidal cells. In one study, *PV*^*+*^ neuron activation changed pyramidal neurons’ firing rates divisively, affecting response gain but not tuning^16^. Another study reported that PVA subtractively lowers pyramidal neuron responses, sharpening orientation tuning^15.^ A third study reported mixed divisive/multiplicative and subtractive/additive effects, which affect response gain but only minimally tuning^17^. Lee et al.^18^ attributed the discrepancy across studies to differences in optogenetic activation parameters^18^, noting that weak or brief *PV*^+^ activation (as in ^16,17^) poorly affected tuning, whereas strong, sustained activation (as in^15^) produced robust sharpening. In contrast to these previous studies, which used narrower irradiance ranges, which also differed across studies, we used a broad range of irradiance levels, leading to changes in firing rate ranging from 0% to −100% (for PVA) and 0% to >+200% (for PVI). This allowed us to observe the full range of effect magnitude seen in the previous mouse studies. Our results partly support the argument by Lee et al.^18^, by demonstrating that at the firing rate changes evoked by the PV manipulation in the studies by^16,17^, the effects on tuning sharpness are much smaller than those seen at the larger firing rate changes caused by PV manipulation in the study by Lee et al^15^. However, Atallah et al.^36^ argued that all the effects observed in these previous mouse studies can be explained by a threshold linear model (TLM) with both multiplicative/divisive and additive/subtractive components, and are consistent with the view that *PV*^*+*^ cells modulate gain, but lead to small changes in tuning at high *PV*^*+*^ neuron activation levels (as observed by^15^), due to the “iceberg effect”^14^. The TLM predicts no changes in tuning for *PV*^*+*^ neuron inactivation. Our results only partly support this argument, as the TLM was able to only account for the linear divisive/multiplicative effects we observed in the D/M neuronal population, but failed to account for the non-linear changes in tuning that we observed in the NL cell population. In particular, we showed that the non-linear sharpening of tuning that we observed in the data cannot be explained by the iceberg effect, for the following 3 reasons: (1) *PV*^*+*^ neuron inactivation (which is not affected by the iceberg effect) significantly broadened tuning; (2) *PV*^*+*^ neuron activation significantly sharpened tuning even at firing rate decreases <30%, for which the TLM model predicts no changes in tuning sharpness; (3) the TLM model overall predicts smaller changes in tuning sharpness than observed in the data for the NL cell population.

Our study is the first to identify multiple, layer-dependent, effects of *PV*^*+*^ neuron manipulation on feature tuning. The linear D/M effects we observed are similar to those reported in mouse V1, but none of the previous studies in mouse reported the NL effects we have observed here. Differently from previous mouse studies, our study used a broader irradiance range, did not pool results across neuronal populations and layers, and did not limit the recordings to superficial layers. These differences in experimental paradigm and analysis may perhaps explain the failure in previous mouse studies of observing NL effects. Interestingly, however, NL effects similar to the ones reported in our study (Mexican-hat-like) were previously observed in cat striate cortex following application of the GABA antagonist bicuculline^1^, although no systematic or quantitative analyses of those effects were performed in that study. Therefore, it remains an open question whether these NL effects are a unique feature of cortical *PV*^+^ neurons in primates and carnivores, species in which sharp orientation tuning uniquely arises in V1, and outside the V1 thalamic input layer in the primate.

## Limitations of the study

A limitation of our study is that it investigated the role of only one type of inhibitory neuron, therefore it remains to be demonstrated whether other inhibitory neuron types in primate V1 also affect tuning or whether this is a unique properties of *PV*^*+*^ cells. Moreover, several models of orientation tuning have postulated that intracortical local recurrent excitation can also play an important role in enhancing orientation tuning^57,68,69^. Therefore, additional studies are needed to disentangle the role of excitation versus inhibition and of different inhibitory neuron types in the sharpening of orientation tuning in primate V1.

A second limitation of our study is that it did not selectively manipulate individual cortical layers or distinct *PV*^*+*^ neuron subtypes. Therefore, we were unable to assess the role of *PV*^+^ cell subtypes in specific layers in isolation. As a result, it is possible that some of the effects we observed in a specific layer were indirect and originated in upstream layer/s^70^.

This is because we lack the necessary technologies. Viral vectors for selective transduction of *PV*^*+*^ neuron subtypes, and high density optoelectronic neural probes for laminar-restricted photostimulation are primarily designed for mouse studies. There is a pressing need for the development of technologies that allow for cell-type specific and spatially-resolved studies of neural circuits in the primate^31,71,72^, the species closest to humans.

## Resource Availability

### Lead Contact

Alessandra Angelucci; email address

### Materials Availability

All viral vectors used in the present study are available for purchase at Addgene, University of Pennsylvania Penn Vector Core, and Azenta Life Sciences gene synthesis and sequencing services.

### Data and Code Availability

Upon acceptance of the study, the data will be made available upon request to the corresponding author.

## Acknowledgments

We thank Kesi Sainsbury and Dr. Matthew Gielow for histological assistance and Sakhtivel Jayakumar for help with microscope imaging. This work was supported primarily by a grant from the National Institute of Health (NIH) to A.A. (R01 EY031959). Additional support was provided by grants from the NIH (R01 EY026812, BRAIN U01 NS099702), the National Science Foundation (IOS 1755431) and the Mary Boesche Endowed Chair, to A.A.; an unrestricted grant from Research to Prevent Blindness, Inc. and a core grant from the NIH (EY014800) to the Department of Ophthalmology, University of Utah.

## Author Contributions

Conceptualization: A.V., A.A., A.M.C; Investigation: all authors; Data Analysis: A.V.; Modeling: A.V.; Writing-Original Draft: A.V, A.M.C., A.A. Writing–Review/Editing: A.A.; Visualization: A.V., A.A.; Supervision & Funding Acquisition: A.A.

## Supplemental Information

The Supplemental Information includes 8 Supplemental Figures with related figure legends.

## STAR Methods

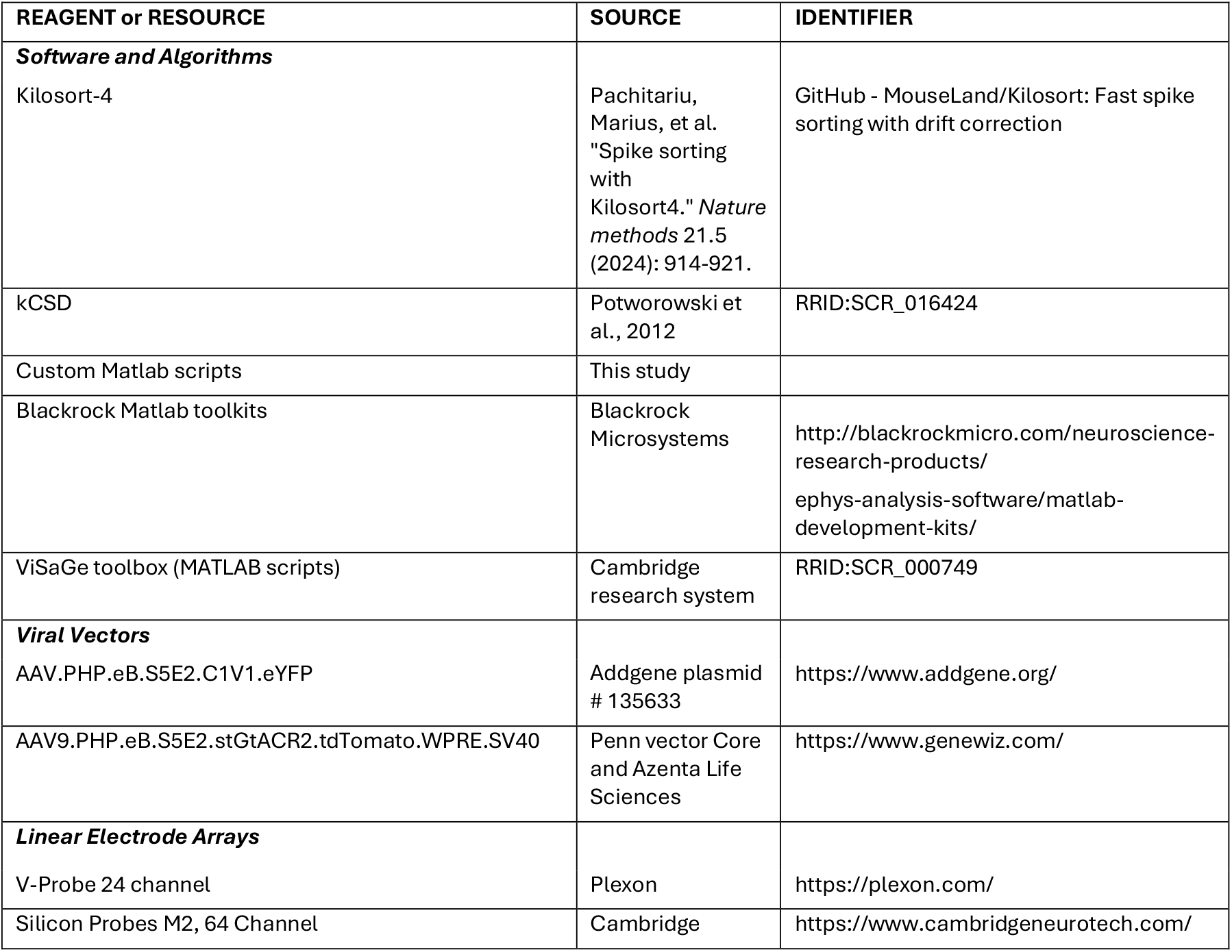

### Animals and Ethics

This study was conducted on four adult common marmosets (*Callithrix jacchus*; two males and two females, 2–8 years old, ~500g body weight) from the University of Utah in-house colony. Electrophysiological recordings were obtained from 11 penetrations in the primary visual cortex (V1) using linear electrode arrays (LEAs). In total, 302 single units (227 orientation-selective, defined as having circular variance, *CV* < 0.96) were recorded under PV-inactivation (PVI) conditions, and 367 single units (310 orientation-selective) under PV-activation (PVA) conditions. All procedures were approved by the University of Utah Institutional Animal Care and Use Committee (IACUC) and adhered to the ethical guidelines outlined by the U.S. Department of Agriculture (USDA) and the National Institutes of Health (NIH).

### Surgical Procedure and Viral Vector Injection

Anesthesia was induced with midazolam (0.1 mg/kg, i.m.) and alfaxalone (10 mg/kg, i.m.), and maintained with isoflurane (1 − 2.5%). An intravenous catheter was inserted into the saphenous vein for continuous infusion of lactated Ringer’s solution (3 − 5ml/kg/hr). Animals were intubated, placed in a stereotaxic apparatus, and artificially ventilated. Physiological parameters (heart rate, oxygen saturation, end-tidal CO_2_, intra-pulmonary pressure, and body temperature) were continuously monitored throughout the experiment. On each hemisphere over area V1, two craniotomies (< 2 mm^2^ each) and small durotomies were made ~-2-4 mm lateral to the midline and ~3-4 mm anterior to the edge of the posterior pole of V1. Each hemisphere received injections of one of two viral vectors: AAV.PHP.eB.S5E2.C1V1.eYFP (2.1E13 GC/ml; Addgene plasmid # 135633) for PVA, or AAV9.PHP.eB.S5E2.stGtACR2.tdTomato.WPRE.SV40 (1.17E13 GC/ml; generation outsourced to the Penn vector Core and Azenta Life Sciences) for PVI. Viral solutions were loaded into glass micropipettes with sharp-beveled tips (~30–45 *μ*m) and slowly pressure-injected (~100–200 nl total) at three depths (1.2, 0.8, and 0.4 mm from the cortical surface) within each craniotomy using a Picospritzer (World Precision Instruments). Following injections, craniotomy sites were sealed with Gelfoam and covered with dental cement. Animals were allowed to recover for 4–6 weeks to ensure optimal viral expression before acute electrophysiological recordings.

### Electrophysiological Recordings

Animals were initially maintained under isoflurane anesthesia (0.5 − 2.5%). Following tracheotomy, a long intravenous catheter (10 cm) was placed in each saphenous vein to facilitate stable anesthesia delivery via continuous infusion. The animal’s head was then secured in a stereotaxic apparatus, the animal was artificially ventilated, and vital signs were continuously monitored as described above. Anesthesia was then transitioned to continuous infusion of sufentanil citrate (4 − 8 *μ*g/kg/hr), and eye movements prevented by continuous infusion of vecuronium bromide (0.3 mg/kg/hr). The pupils were dilated by topical application of atropine, the corneas protected with gas-permeable contact lenses, and the eyes were refracted. The craniotomies were re-opened, the regrown dura removed, and the viral injected sites were identified by reporter protein expression visualized using a fluorescent surgical microscope (Carl Zeiss, GmbH). Laminar extracellular recordings were made using linear electrode arrays (LEAs), either 24-channel V-Probes (Plexon, Dallas, TX; 100µm contact spacing, 20 µm contact diameter) or 64-Channel Cambridge NeuroTech (Cambridge, UK; 31µm intercontact spacing, 11×15µm contact diameter). LEAs were lowered normally to the cortical surface (using triangulation methods) through a custom-made guide tube, with the electrode advanced slowly (1 *μ*m/s) into the cortex to a depth of up to 2.2 mm, guided by monitoring of neuronal signals. The exposed cortical tissue around the guide tube was sealed with agar and Dura-Gel (Cambridge NeuroTech, Cambridge, UK) to prevent desiccation and stabilize the recordings. Recording signals were amplified, digitized and sampled at 30kHz using a 128-channel system (Cerebus, Blackrock Microsystems, Salt Lake City, UT). Real-time online analysis was performed using Blackrock’s MATLAB function *cbmex*. Spikes were detected as spatiotemporal waveforms using the spike sorting algorithm Kilosort4 and manually validated^73^.

### Visual Stimuli

Visual stimuli were generated in MATLAB (Mathworks Inc., Natick, MA; RRID:SCR_001622) and presented on a calibrated CRT monitor (Sony GDM-C520K; 100 Hz refresh rate; resolution: 768 × 1024 pixels; mean luminance: 45 cd/m^2^; viewing distance: 57 cm). Stimulus presentation and synchronization were controlled via a ViSaGe system (Cambridge Research Systems, Cambridge, UK; RRID:SCR_000749). Minimum response fields (mRFs) across contacts were first mapped using sparse noise^74^ and Hartley stimuli^75^. Subsequent stimuli were centered on the aggregate mRF of the column. Drifting sinusoidal grating patches of 100% contrast and varying parameters were then presented monocularly, to determine each unit’s preferred spatial frequency (SF), temporal frequency (TF), and size. Finally, orientation-tuning functions across the LEA were measured using drifting grating patches at the preferred SF, TF, and size for most contacts across the LEA varying in drift direction (24 or 36 directions; 5–10 repeats per direction; 1 s stimulus duration; 2 s inter-stimulus interval), with and without laser photostimulation. To monitor eye movements, mRFs were re-mapped approximately every 10-20 min.

### Laser Photostimulation

Surface photostimulation at the LEA recording sites was delivered using either a blue laser (wavelength: *473* nm) or a green laser (wavelength: *532* nm) coupled to a 400 µm-diameter optical fiber and collimator lens, yielding a final spot size measured on the cortical surface of *1*.*3* mm^2^. Because the laser beam traveled through agar and cortical tissue, direct optical measurements could not accurately estimate the effective light power at the target site. Moreover, potential variability in viral expression between animals and across penetration sites precluded using laser output power as a reliable measure of stimulation strength. Instead, we quantified laser efficacy by calculating the percent change in population firing rate (ΔFr) during combined visual and laser stimulation relative to the visual stimulation-only condition. This measure was used to parameterize the functional “laser power” for analysis. To sample a broad range of manipulations (0 to −100% ΔFr for PVA; 0 to > 200% ΔFr for PVI), we tested, in separate experiments, multiple laser powers per penetration (irradiance range: 0.1-0.3mW/mm^2^ for PVI, and 0.3-1.8mW/mm^2^ for PVA; measured as laser power exciting the collimator, divided by the area of the collimator).

### Quantitative Data Analysis

#### Neuronal sample selection and clustering

We recorded from a total of 669 spike-sorted single units across 11 penetrations in 4 animals. From these units, we excluded from analysis non-orientation selective units (*CV* > 0.96; n = 132 cells), and opsin-tagged *PV*^+^ units (n=42), i.e., those showing increased (decreased) firing rate during PVA (PVI).

The remaining units (n=495) were clustered using the second and fourth derivatives of the *ΔT* curves (difference between tuning curves; **Fig. 1B-C**, fourth row, *red curves*) computed at the neuron’s preferred orientation (see **Fig. S2A-C**). Units with an upward peak at the preferred orientation (negative second derivative), or with a flat curve at the preferred orientation (both positive second and fourth derivatives) were classified as *non-linear* (NL, *yellow dots* in **Fig. S2A**). For the remaining units, i.e. those with a downward peak at the preferred orientation in the *ΔT* curves (negative fourth derivative), we computed a multiplicative ratio:

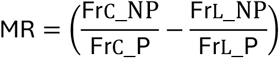

where Fr_C__NP and Fr_L__NP are the control (no-laser) and laser firing rates at the non-preferred (NP) orientation, and Fr_C__P and Fr_L__P are the control and laser firing rates at the preferred orientation, respectively. Preferred and non-preferred orientations were defined as the orientation at max and min response in the fitted tuning curves, respectively (as detailed below). Units with MR < 0.02 were classified as *linear multiplicative/divisive* (D/M, *black dots* in **Fig. S2A**). The *RT* curves (ratio between laser and control tuning curves) for these units were flat horizontal lines (**Fig. 1B** fourth row, and **Fig. 1D** bottom row, *cyan curves*; **Fig. S2C** Left column, bottom 2 panels). To quantify the “flatness” of the *RT* curve, we measured the Coefficient of Variation (CoV) as the standard deviation (SD)/mean of the *RT* curve. The CoV for the D/M population was significantly smaller (mean CoV±s.e.m, 0.091±0.0058, p=2.5×10^−32^, PVA; 0.049±0.0058, p=2.8×10^−15^, PVI, Wilcoxon rank-sum test) than the CoV for the NL population (0.353±0.0187, PVA; 0.143±0.009, PVI), indicating the *RT* curve for the D/M population indeed approximates a flat horizontal line (**Fig. S2D**). We also computed the CoV for the Δ*T* curves, and found that both the D/M and NL populations showed large CoV values (mean±s.e.m, D/M: 0.432±0.0183, p=0.0089, PVA; 0.251±0.0253, p=5.8×10^−5^ PVI; NL: 0.377±0.0148, PVA; 0.177±0.0137, PVI) indicative of lack of subtractive/additive effects of *PV*^*+*^ cell manipulations (**Fig. S2D**).

Units that could not be classified as NL or D/M based on the above criteria, *red and white dots* in **Fig. S2A**, and **Fig. S2C** right column) were unclassified (U). The Unclassified population comprised a small population of units (Mix, *white dots* in **Fig. S2A**) that showed mixed D/M and NL effects at different laser intensities, and cells that could not be categorized (Uct, *red dots* in **Fig. S2A**), into any of these groups (see Results for details). All 495 units were included in the population analyses; however, for many of the analyses (those reported in **Figs. 2,3,4A-B,5,8,S2A,C,D, S5A-B, S6, S7, S8**) the number of units reported is larger, because the same units recorded at different laser intensities were considered independent samples. Importantly, we matched waveforms across laser intensities and verified that units classified as D/M or NL showed the same effect at all intensities. Those that did not satisfy this criterion were considered unclassified (n=29, Mix group).

#### Statistical model fitting and computations of orientation tuning metrics

Orientation-tuning data were fitted with von Mises functions^76^ of the form:

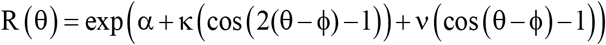

where *θ* is stimulus orientation (in degrees), *ϕ* is the central orientation of the tuning, *κ* weights the second harmonic (cos 2Δ), *ν* weights the first harmonic (cos Δ), and *α* is a log-baseline/gain term. Because the model includes both first- and second-harmonic terms, the distance between the first and second peaks is not fixed at 180°, providing the flexibility to capture asymmetric tuning commonly observed in neural data and yielding better fits than models that enforce a strict 180° separation. Accordingly, we define the preferred orientation (θ_P) as the orientation at which R(θ) attains its global maximum, and the non-preferred orientation (θ_NP) as the orientation at which R(θ) attains its minimum. In our dataset, θ_NP lies very close to the orthogonal of θ_pref (≈90° apart), consistent with the empirical distribution of angular distances between the maximum and minimum responses (see **Fig. S1B**).

From the fitted functions we extracted the following several metrics of tuning:

The orientation selectivity index (OSI) as:

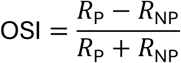

where R_P_ is the response to the preferred orientation, and R_NP_ is the response to the non-preferred orientation defined as described above.

The half-bandwidth (HBW), defined as half the difference between the two orientation values on either side of the function’s peak that correspond to the half-maximum response level measured as:

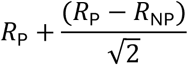

The circular variance (CV)^25^, computed as:

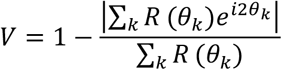

where *R*(*θ*_*k*_) is the response magnitude of the neuron to stimulus orientation *θ*_*k*_; each sampled orientation *θ*_*k*_ is treated as unit vector on the complex plane at angle 2*θ*_;_and weighed by *R*(*θ*_*k*_). Summing these weighted vectors gives a resultant vector, whose magnitude (after dividing by the total weight ∑_*k*_ *R*(*θ*_*k*_)) is the resultant length. CV is 1minus that length, and has a value of 1 for a dataset falling uniformly on a circle (non-orientation selective), and a value of 0 for a dataset with response only at a single orientation (highly selective).

For OSI, HBW, CV, and firing rate (Fr), the delta (Δ) was defined as the difference between the values in the laser and control conditions.

#### Laminar border identification

We used multiple criteria to assign units to specific layers, based on both stimulus-evoked and spontaneous MUA and SUA, as well as stimulus-evoked current source density (CSD^77^) and spectral analyses (**Fig. S4**). For CSD analysis we followed the methods of Bijanzadeh et al.^78^. Specifically, signals were band-pass filtered (1–100 Hz), the stimulus-evoked LFPs were trial averaged, the second spatial derivative (kernel-CSD) was estimated, and baseline corrected (z-scored) to visualize current sinks (negative deflections) and sources (positive deflections; **Fig. S4D**). We additionally computed stimulus-evoked local coherence spectra as in^79^ (**Fig. S4E**); specifically, for each contact we estimated the magnitude-squared coherence with its immediately neighboring contacts across different frequencies, and baseline corrected the stimulus-evoked coherence.

Using these analyses, we identified the top of the cortex as one contact above the most superficial contact showing spiking activity (in the example of **Fig. S4A** this was contact #1), to account for layer 1. The layer 6/white matter boundary was identified based on a combination of cortical depth (range of 1.6-1.9mm from top contact) and spiking profile (a drop in visually-driven activity, and increased spontaneous activity). The G layer was identified as a 300-350µm-thick band around the middle of the cortical depth showing: (1) highest spontaneous and visually-driven activity (**Fig. S4A,B**), (2) shortest inter-spike (ISI) interval (**Fig. S4C**), (3) Strong coherence (**Fig. S4E**), and (4) the earliest current sink in the CSD profile (**Fig. S4D**). The G layer boundaries were particularly evident in the spectral coherence analysis, and the lower boundary of the G layer in the coherence corresponded to the border between the earliest current sink and a reversal to source in the CSD, as previously shown^80^. Units above and below the G layer were assigned to the SG and IG layers, respectively.

### Statistical Analyses

Statistical analyses were performed on orientation-tuning curves fit with von Mises– functions, from which standard selectivity metrics (OSI, CV and HBW)—were computed. For all measures, laser effects were summarized as Δ (laser minus control) values. Linear regression was used to quantify the relationships between the laser-induced firing-rate change (ΔFr) and changes in tuning metrics (e.g., OSI, CV, HBW), and the significance of these regressions was evaluated across the population (**Figs. 2-3,8, S6,S7, S8**). For within-unit paired comparisons between control and laser conditions we employed the Wilcoxon signed-rank test (**Figs. 2,3,5**), while for between-group comparisons we used the non-parametric Wilcoxon rank-sum tests (**Fig. 4**), as appropriate. The specific tests used for each analysis are detailed in the Results and/or Figure Legends.

### Post-mortem Histology

On completion of the recordings, animals were sacrificed with Beuthanasia (0.22ml/kg, i.p.) and perfused with 4% paraformaldehyde in 0.1M phosphate buffer for 20 minutes. The blocks containing V1 were frozen-sectioned at 40µm, sagittally. Reporter protein expression was visualized under fluorescent microscopy to validate viral expression in V1 (**Fig. 1A**), and appropriate targeting of LEA penetrations to regions of viral expression.

### Computational Models

To evaluate the predictive accuracy of the threshold linear model (TLM), we simulated laser tuning curves by applying a pure linear threshold transformation to the control tuning curves as in^17^. First, we fitted the model (including a threshold for PVA) using the control tuning curve as the independent variable (x-axis) and the corresponding laser tuning curve as the dependent variable (y-axis; as in **Fig. 1B** fifth column). This fitting yielded two parameters: slope and offset. Using these parameters, we simulated laser tuning curves by applying the TLM to the control curve. This procedure enabled us to estimate the expected laser response under the model’s assumptions. The simulated curves were then compared to the actual laser tuning curves to quantify the model’s accuracy, by regression and estimation of the R-squared as a metric of goodness-of-fit. The same procedure was also applied to our I/O model. For each model, we then compared the simulated and actual laser tuning curves by examining changes in orientation tuning (HBW, OSI and CV), as a function of the relative change in firing rate:

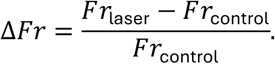

The extent to which each model reproduced the empirical relationships (**Figs. S6-S8, 8**) was used as a measure of its explanatory power.

The I/O model is described in the Results.

## References

1. Sillito, A.M. (1979). inhibitory mechanisms influencing complex cell orientation selectivity and their modification at high resting discharge levels. J. Physiol. (Lond.) 289, 33–53.

2. Katzner, S., Busse, L., and Carandini, M. (2011). GABAA inhibition controls response gain in visual cortex. J. Neurosci. 31, 5931–5941. 10.1523/JNEUROSCI.5753-10.2011.

3. Isaacson, J.S., and Scanziani, M. (2011). How inhibition shapes cortical activity. Neuron 72, 231–243.

4. Zhu, Y., Qiao, W., Liu, K., Zhong, H., and Yao, H. (2015). Control of response reliability by parvalbumin-expressing interneurons in visual cortex. Nat. Commun. 6, 6802. 10.1038/ncomms7802.

5. Sohal, V.S., Zhang, F., Yizhar, O., and Deisseroth, K. (2009). Parvalbumin neurons and gamma rhythms enhance cortical circuit performance. Nature 459, 698–702. 10.1038/nature07991.

6. Chen, A., Sun, Y., Lei, Y., Li, C., Liao, S., Meng, J., Bai, Y., Liu, Z., Liang, Z., Zhu, Z., et al. (2023). Single-cell spatial transcriptome reveals cell-type organization in the macaque cortex. Cell 186, 3726–3743 e3724. 10.1016/j.cell.2023.06.009.

7. DeFelipe, J. (1997). Types of neurons, synaptic connections and chemical characteristics of cells immunoreactive for calbindin-D28K, parvalbumine and calretinin in the neocortex. J. Chem. Neuroanat. 14, 1–19.

8. Somogyi, P., Tamas, G., Lujan, R., and Buhl, E.H. (1998). Salient features of synaptic organisation in the cerebral cortex. Brain. Res. Brain. Res. Rev. 26, 113–135.

9. Kawaguchi, Y., and Kubota, Y. (1997). GABAergic cell subtypes and their synaptic connections in rat frontal cortex. Cereb. Cortex 7, 476–486.

10. Tremblay, R., Lee, S., and Rudy, B. (2016). GABAergic Interneurons in the Neocortex: From Cellular Properties to Circuits. Neuron 91, 260–292.

11. Hubel, D.H., and Wiesel, T.N. (1959). Receptive fields of single neurones in the cat’s striate cortex. J. Physiol. 148, 574–591.

12. Ferster, D., and Miller, K.D. (2000). Neural mechanisms of orientation selectivity in the visual cortex. Ann. Rev. Neurosci. 23, 441–471.

13. Sompolinsky, H., and Shapley, R. (1997). New perspectives on the mechanisms for orientation selectivity. Current opinion in neurobiology 7, 514–522.

14. Priebe, N.J., and Ferster, D. (2008). Inhibition, spike threshold, and stimulus selectivity in primary visual cortex. Neuron 57, 482–497.

15. Lee, S.H., Kwan, A.C., Zhang, S., Phoumthipphavong, V., Flannery, J.G., Masmanidis, S.C., Taniguchi, H., Huang, Z.J., Zhang, F., Boyden, E.S., et al. (2012). Activation of specific interneurons improves V1 feature selectivity and visual perception. Nature 488, 379–383.

16. Wilson, N.R., Runyan, C.A., Wang, F.L., and Sur, M. (2012). Division and subtraction by distinct cortical inhibitory networks in vivo. Nature 488, 343–348.

17. Atallah, B.V., Bruns, W., Carandini, M., and Scanziani, M. (2012). Parvalbumin-expressing interneurons linearly transform cortical responses to visual stimuli. Neuron 73, 159–170.

18. Lee, S.H., Kwan, A.C., and Dan, Y. (2014). Interneuron subtypes and orientation tuning. Nature 508, E1–2.

19. Piscopo, D.M., El-Danaf, R.N., Huberman, A.D., and Niell, C.M. (2013). Diverse visual features encoded in mouse lateral geniculate nucleus. J. Neurosci. 33, 4642–4656.

20. Goetz, J., Jessen, Z.F., Jacobi, A., Mani, A., Cooler, S., Greer, D., Kadri, S., Segal, J., Shekhar, K., Sanes, J.R., and Schwartz, G.W. (2022). Unified classification of mouse retinal ganglion cells using function, morphology, and gene expression. Cell Rep. 40, 111040. 10.1016/j.celrep.2022.111040.

21. Sun, W., Tan, Z., Mensh, B.D., and Ji, N. (2016). Thalamus provides layer 4 of primary visual cortex with orientation- and direction-tuned inputs. Nat. Neurosci. 19, 308–315. 10.1038/nn.4196.

22. Scholl, B., Tan, A.Y., Corey, J., and Priebe, N.J. (2013). Emergence of orientation selectivity in the Mammalian visual pathway. J. Neurosci. 33, 10616–10624.

23. Zhao, X., Chen, H., Liu, X., and Cang, J. (2013). Orientation-selective responses in the mouse lateral geniculate nucleus. J. Neurosci. 33, 12751–12763.

24. Wang, H., Dey, O., Lagos, W.N., Behnam, N., Callaway, E.M., and Stafford, B.K. (2024). Parallel pathways carrying direction-and orientation-selective retinal signals to layer 4 of the mouse visual cortex. Cell Rep. 43, 113830. 10.1016/j.celrep.2024.113830.

25. Ringach, D.L., Shapley, R.M., and Hawken, M.J. (2002). Orientation selectivity in macaque V1: diversity and laminar dependence. J. Neurosci. 22, 5639–5651.

26. Hawken, M.J., and Parker, A. (1984). Contrast sensitivity and orientation selectivity in laminar IV of the striate cortex of old world monkeys. Exp. Brain Res. 54, 367–372.

27. Wang, T., Li, Y., Yang, G., Dai, W., Yang, Y., Han, C., Wang, X., Zhang, Y., and Xing, D. (2020). Laminar Subnetworks of Response Suppression in Macaque Primary Visual Cortex. J. Neurosci. 40, 7436–7450. 10.1523/JNEUROSCI.1129-20.2020.

28. Goodchild, A.K., and Martin, P.R. (1998). The distribution of calcium-binding proteins in the lateral geniculate nucleus and visual cortex of a New World monkey, the marmoset, Callithrix jacchus. Vis. Neurosci. 15, 625–642.

29. DeFelipe, J., Gonzalez-Albo, M.C., Del Rio, M.R., and Elston, G.N. (1999). Distribution and patterns of connectivity of interneurons containing calbindin, calretinin, and parvalbumin in visual areas of the occipital and temporal lobes of the macaque monkey. J. Comp. Neurol. 412, 515–526.

30. Xu, X., Roby, K.D., and Callaway, E.M. (2010). Immunochemical characterization of inhibitory mouse cortical neurons: three chemically distinct classes of inhibitory cells. J Comp Neurol 518, 389–404. 10.1002/cne.22229.

31. Federer, F., Balsor, J., Ingold, A., Babcock, D.P., Dimidschstein, J., and Angelucci, A. (2024). Laminar specificity and coverage of viral-mediated gene expression restricted to GABAergic interneurons and their parvalbumin subclass in marmoset primary visual cortex. eLife 13, RP97673.

32. Medalla, M., Mo, B., Nasar, R., Zhou, Y., Park, J., and Luebke, J.I. (2023). Comparative features of calretinin, calbindin, and parvalbumin expressing interneurons in mouse and monkey primary visual and frontal cortices. J. Comp. Neurol. 531, 1934–1962. 10.1002/cne.25514.

33. Prakash, R., Yizhar, O., Grewe, B., Ramakrishnan, C., Wang, N., Goshen, I., Packer, A.M., Peterka, D.S., Yuste, R., Schnitzer, M.J., and Deisseroth, K. (2012). Two-photon optogenetic toolbox for fast inhibition, excitation and bistable modulation. Nat. Methods 9, 1171–1179. 10.1038/nmeth.2215.

34. Mahn, M., Gibor, L., Patil, P., Cohen-Kashi Malina, K., Oring, S., Printz, Y., Levy, R., Lampl, I., and Yizhar, O. (2018). High-efficiency optogenetic silencing with soma-targeted anion-conducting channelrhodopsins. Nat. Commun. 9, 4125. 10.1038/s41467-018-06511-8.

35. Vormstein-Schneider, D.C., Lin, J.D., Pelkey, K.A., Chittajallu, R., Guo, B., Arias Garcia, M., Sakopoulos, S., Stevenson, O., Schneider, G., Zhang, Q., et al. (2020). Viral manipulation of functionally distinct interneurons in mice, non-human primates and humans. Nat. Neurosci. 23, 1629–1636. 10.1101/808170.

36. Atallah, B.V., Scanziani, M., and Carandini, M. (2014). Atallah et al. reply. Nature 508, E3. 10.1038/nature13129.

37. Shapley, R., Hawken, M., and Ringach, D.L. (2003). Dynamics of orientation selectivity in the primary visual cortex and the importance of cortical inhibition. Neuron 38, 689–699. 10.1016/s0896-6273(03)00332-5.

38. Carandini, M., and Ferster, D. (2000). Membrane potential and firing rate in cat primary visual cortex. J. Neurosci. 20, 470–484. 10.1523/JNEUROSCI.20-01-00470.2000.

39. Ferguson, K.A., and Cardin, J.A. (2020). Mechanisms underlying gain modulation in the cortex. Nat. Rev. Neurosci. 21, 80–92. 10.1038/s41583-019-0253-y.

40. Gerstner, W., Kistler, W.M., Naud, R., and Paninski, L. (2014). Neuronal dynamics: From single neurons to networks and models of cognition (Cambridge University Press).

41. Finn, I.M., Priebe, N.J., and Ferster, D. (2007). The emergence of contrast-invariant orientation tuning in simple cells of cat visual cortex. Neuron 54, 137–152. 10.1016/j.neuron.2007.02.029.

42. Chance, F.S., Abbott, L.F., and Reyes, A.D. (2002). Gain modulation from background synaptic input. Neuron 35, 773–782. 10.1016/s0896-6273(02)00820-6.

43. Anderson, J.S., Lampl, I., Gillespie, D.C., and Ferster, D. (2000). The contribution of noise to contrast invariance of orientation tuning in cat visual cortex. Science 290, 1968–1972. 10.1126/science.290.5498.1968.

44. Miller, K.D., and Troyer, T.W. (2002). Neural noise can explain expansive, power-law nonlinearities in neural response functions. J. Neurophysiol. 87, 653–659. 10.1152/jn.00425.2001.

45. Hansel, D., and van Vreeswijk, C. (2002). How noise contributes to contrast invariance of orientation tuning in cat visual cortex. J. Neurosci. 22, 5118–5128. 10.1523/JNEUROSCI.22-12-05118.2002.

46. Mensi, S., Naud, R., and Gerstner, W. (2011). From Stochastic Nonlinear Integrate-and-Fire to Generalized Linear Models.

47. Pillow, J.W., Shlens, J., Paninski, L., Sher, A., Litke, A.M., Chichilnisky, E.J., and Simoncelli, E.P. (2008). Spatio-temporal correlations and visual signalling in a complete neuronal population. Nature 454, 995–999. 10.1038/nature07140.

48. Rullan Buxo, C.E., and Pillow, J.W. (2020). Poisson balanced spiking networks. PLoS Comput. Biol. 16, e1008261. 10.1371/journal.pcbi.1008261.

49. Azouz, R. (2005). Dynamic spatiotemporal synaptic integration in cortical neurons: neuronal gain, revisited. J. Neurophysiol. 94, 2785–2796. 10.1152/jn.00542.2005.

50. Higgs, M.H., Slee, S.J., and Spain, W.J. (2006). Diversity of gain modulation by noise in neocortical neurons: regulation by the slow afterhyperpolarization conductance. J. Neurosci. 26, 8787–8799. 10.1523/JNEUROSCI.1792-06.2006.

51. Vafaei, A., Mohammadi, M., Khadir, A., Zabeh, E., YazdaniBanafsheDaragh, F., Khorasani, M., and Lashgari, R. (2021). V1 receptive field structure contributes to neuronal response latency. bioRxiv DOI: 10.1101/2021.12.30.474591.

52. Krienen, F.M., Levandowski, K.M., Zaniewski, H., Del Rosario, R.C.H., Schroeder, M.E., Goldman, M., Wienisch, M., Lutservitz, A., Beja-Glasser, V.F., Chen, C., et al. (2023). A marmoset brain cell census reveals regional specialization of cellular identities. Sci. Adv. 9, eadk3986. 10.1126/sciadv.adk3986.

53. Blumcke, I., Hof, P.R., Morrison, J.H., and Celio, M.R. (1990). Distribution of parvalbumin immunoreactivity in the visual cortex of Old World monkeys and humans. J Comp Neurol 301, 417–432. 10.1002/cne.903010307.

54. Lund, J.S., and Yoshioka, T. (1991). Local circuit neurons of macaque monkey striate cortex: III. Neurons of laminae 4B, 4A and 3B. J. Comp. Neurol. 331, 234–258.

55. Lund, J.S., and Wu, C.Q. (1997). Local circuit neurons of macaque monkey striate cortex: IV. Neurons of laminae 1-3A. J. Comp. Neurol. 384, 109–126.

56. Ben-Yishai, R., Bar-Or, R.L., and Sompolinsky, H. (1995). Theory of orientation tuning in visual cortex. Proc. Natl. Acad. Sci. USA 92, 3844–3848.

57. Somers, D.C., Nelson, S.B., and Sur, M. (1995). An emergent model of orientation selectivity in cat visual cortical simple cells. J. Neurosci. 15, 5448–5465.

58. Lund, J.S., Angelucci, A., and Bressloff, P.C. (2003). Anatomical substrates for functional columns in macaque monkey primary visual cortex. Cereb. Cortex 13, 15–24.

59. Lund, J.S. (1987). Local circuit neurons of macaque monkey striate cortex: I. Neurons of laminae 4C and 5A. J. Comp. Neurol. 257, 60–92. 10.1002/cne.902570106.

60. Kerlin, A.M., Andermann, M.L., Berezovskii, V.K., and Reid, R.C. (2010). Broadly tuned response properties of diverse inhibitory neuron subtypes in mouse visual cortex. Neuron 67, 858–871.

61. Ma, W.P., Liu, B.H., Li, Y.T., Huang, Z.J., Zhang, L.I., and Tao, H.W. (2010). Visual representations by cortical somatostatin inhibitory neurons--selective but with weak and delayed responses. J. Neurosci. 30, 14371–14379.

62. Dai, W., Clark, A.G., Vafaei, A., Federer, F., and Angelucci, A. (2025). Functional tuning of parvalbumin-expressing interneurons across the layers of primate primary visual cortex. Soc. Neurosci. Abstr. Online.

63. Stringer, C., Pachitariu, M., Steinmetz, N.A., Okun, M., Bartho, P., Harris, K.D., Sahani, M., and Lesica, N.A. (2016). Inhibitory control of correlated intrinsic variability in cortical networks. Elife 5, e19695. 10.7554/eLife.19695.

64. Hennequin, G., Ahmadian, Y., Rubin, D.B., Lengyel, M., and Miller, K.D. (2018). The Dynamical Regime of Sensory Cortex: Stable Dynamics around a Single Stimulus-Tuned Attractor Account for Patterns of Noise Variability. Neuron 98, 846–860 e845. 10.1016/j.neuron.2018.04.017.

65. Sadeh, S., and Clopath, C. (2021). Inhibitory stabilization and cortical computation. Nat. Rev. Neurosci. 22, 21–37. 10.1038/s41583-020-00390-z.

66. Bernacchia, A., and Wang, X.J. (2013). Decorrelation by recurrent inhibition in heterogeneous neural circuits. Neural Comput. 25, 1732–1767. 10.1162/NECO_a_00451.

67. Sadagopan, S., and Ferster, D. (2012). Feedforward origins of response variability underlying contrast invariant orientation tuning in cat visual cortex. Neuron 74, 911–923. 10.1016/j.neuron.2012.05.007.

68. Hansel, D., and Sompolinsky, H. (1996). Chaos and synchrony in a model of a hypercolumn in visual cortex. J. Comput. Neurosci. 3, 7–34. 10.1007/BF00158335.

69. Koch, E., Jin, J., Alonso, J.M., and Zaidi, Q. (2016). Functional implications of orientation maps in primary visual cortex. Nat. Commun. 7, 13529. 10.1038/ncomms13529.

70. Moore, A.K., Weible, A.P., Balmer, T.S., Trussell, L.O., and Wehr, M. (2018). Rapid Rebalancing of Excitation and Inhibition by Cortical Circuitry. Neuron 97, 1341–1355 e1346. 10.1016/j.neuron.2018.01.045.

71. Clark, A.M., Ingold, A., Reiche, C.F., Cundy, D., 3rd, Balsor, J.L., Federer, F., McAlinden, N., Cheng, Y., Rolston, J.D., Rieth, L., et al. (2024). An optrode array for spatiotemporally-precise large-scale optogenetic stimulation of deep cortical layers in non-human primates. Commun. Biol. 7, 329. 10.1038/s42003-024-05984-2.

72. Gielow, M., Federer, F., Jayakumar, A., Caldwell, Z., Winebrenner, C., Xu, X., and Angelucci, A. (2025). Laminar specificity and coverage of enhancer-AAV-mediated gene expression restricted to subpopulations of Somatostatin and Parvalbumin inhibitory interneurons in primate V1. Soc. Neurosci. Abstr. Online.

73. Pachitariu, M., Sridhar, S., Pennington, J., and Stringer, C. (2024). Spike sorting with Kilosort4. Nat. Methods 21, 914–921. 10.1038/s41592-024-02232-7.

74. Jones, J.P., and Palmer, L.A. (1987). The two-dimensional spatial structure of simple receptive fields in cat striate cortex. J. Neurophysiol. 58, 1187–1211.

75. Ringach, D.L., Sapiro, G., and Shapley, R. (1997). A subspace reverse-correlation technique for the study of visual neurons. Vision Res. 37, 2455–2464. 10.1016/s0042-6989(96)00247-7.

76. Ringach, D.L., Hawken, M.J., and Shapley, R. (2003). Dynamics of orientation tuning in macaque V1: the role of global and tuned suppression. J. Neurophysiol. 90, 342–352. 10.1152/jn.01018.2002.

77. Mitzdorf, U. (1985). Current source-density method and application in cat cerebral cortex: investigation of evoked potentials and EEG phenomena. Physiol Rev 65, 37–100.

78. Bijanzadeh, M., Nurminen, L., Merlin, S., and Angelucci, A. (2018). Distinct laminar processing of local and global context in primate primary visual cortex. Neuron 100, 1–16.

79. Zhang, L.A., Li, P., and Callaway, E.M. (2024). High-Resolution Laminar Identification in Macaque Primary Visual Cortex Using Neuropixels Probes. bioRxiv. 10.1101/2024.01.23.576944.

80. Schroeder, C.E., Mehta, A.D., and Givre, S.J. (1998). A spatiotemporal profile of visual system activation revealed by current source density analysis in the awake macaque. Cereb. Cortex 8, 575–592.

