## Supplemental Figs 1-8 for "Laminar-specific control of response gain and orientation-tuning by parvalbumin-expressing inhibitory interneurons in primate visual cortex"

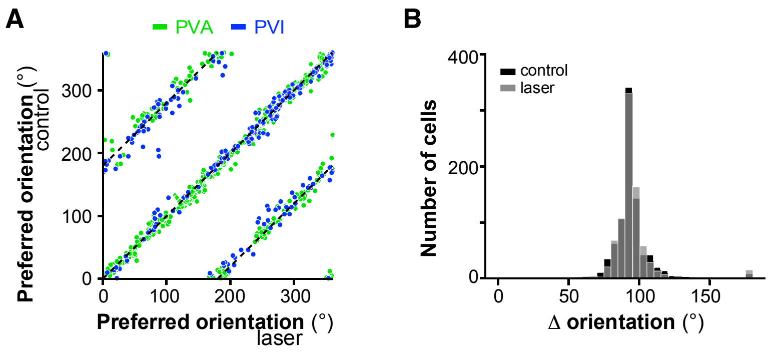


**Figure S1** (Related to Figs. 1,3)

***PV***$^{+}$ **interneuron manipulation does not affect the preferred orientation. (A)** The preferred orientation in the control condition is plotted against the preferred orientation in the laser condition, for cells recorded in PVA (*green*) and PVI (*blue*) experiments. Cells along the diagonal line do not change their preferred orientation. Cells along the lines with slope of 1 switched their preferred direction (± 180°) but not orientation. **(B)** Distribution of angular distance between the preferred and non-preferred orientation across the cell population, in the control (*black*) and laser (*gray*) conditions.

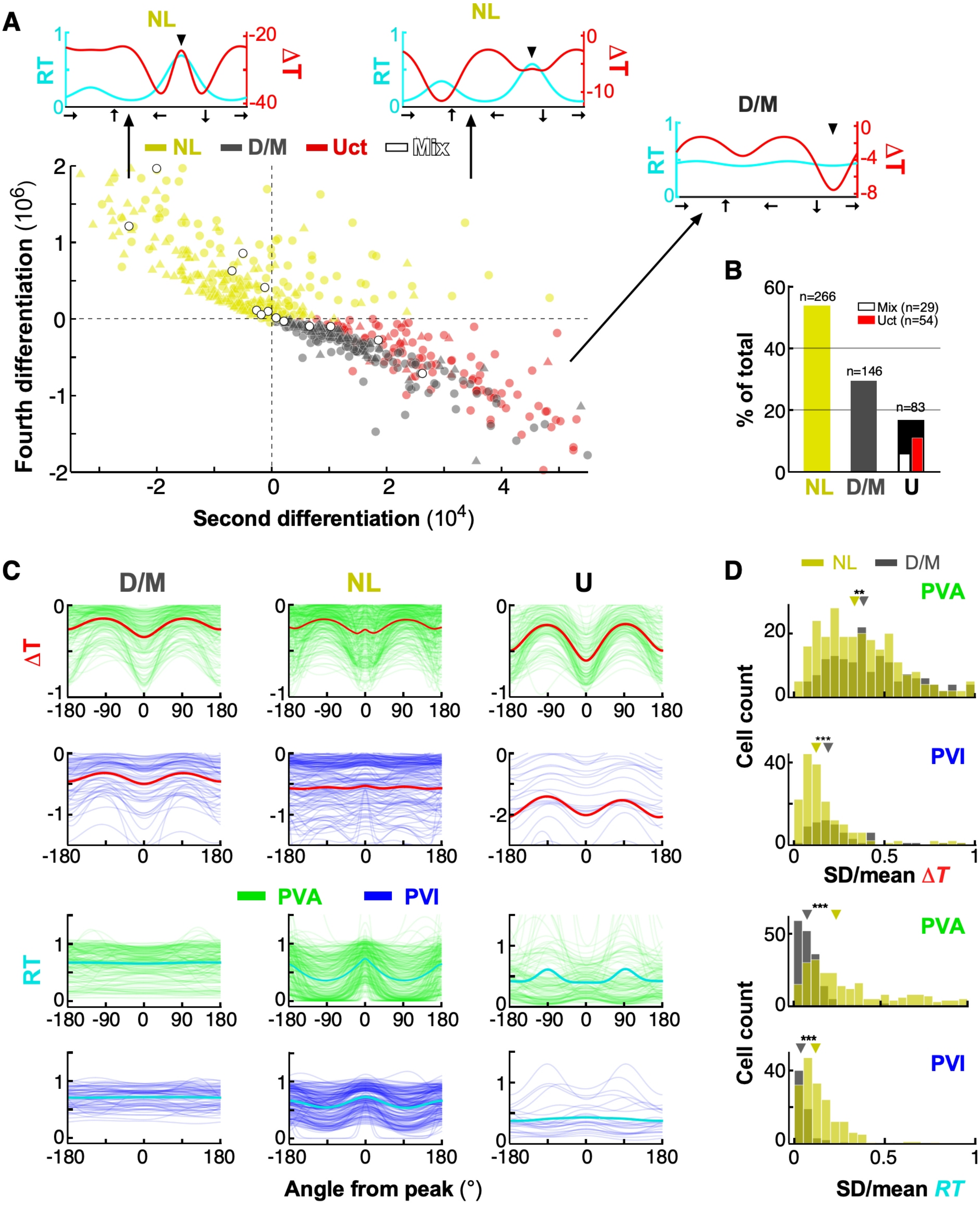


**Figure S2** (Related to Fig. 1)

**Clustering of recorded units.**
**(A)** Scatter plot of all recorded units in a 2D feature space, where the x-axis represents the second derivative, and the Y-axis the fourth derivative of the $\Delta T$ curve, computed at the curve's peak (see STAR Methods). *Yellow dots:* units classified as NL; *gray dots*: units classified as linear D/M; *red dots*: uncategorized units (Uct); *white dots:* mixed units (Mix). *Circles and triangles:* units recorded under PVA and PVI experiments, respectively. *Insets:* representative $\Delta T$ and *RT* curves in each quadrant of the 2D feature space. *Arrowheads* mark the cell's preferred orientation. **(B)** Percent of units falling into each group. Here each unit recorded at different light intensities is considered a single sample, while in (A) units recorded at different intensities are considered independent samples. Units classified as NL (*yellow bar*) or D/M (*gray bar*) showed the same effect at all intensities. Mix unclassified cells (*white bar*) showed D/M or NL effects at different laser intensities. *Black bar:* unclassified units that did not fit into any category (Uct). n= number of units in each group. **(C)** Top two rows: $\Delta T$ curves (computed from normalized tuning curves) for each individual unit under PVA (top row, *green*) and PVI (second row, *blue*) experiments, grouped by effect type (Left: D/M; Middle: NL; Right: U). *Red curves*: population averages. Bottom two rows: Same as Top two rows, but here *RT* curves are shown, instead, and *cyan curves* are population averages. **(D)** Top two rows: distribution of Coefficient of Variation (CoV=standard deviation/mean) of the $\Delta T$ curve across the population of NL and D/M units under PVA (top row) and PVI (second row) experiments. Most cells in both populations show large CoV values indicating lack of subtractive/additive effects of $PV^+$ cell manipulation. Bottom two rows: distribution of CoV of the *RT* curve for the NL and D/M populations under PVA (top row) and PVI (second row). The D/M population shows significantly smaller CoV thatn the NL population, indicative of D/M effects.

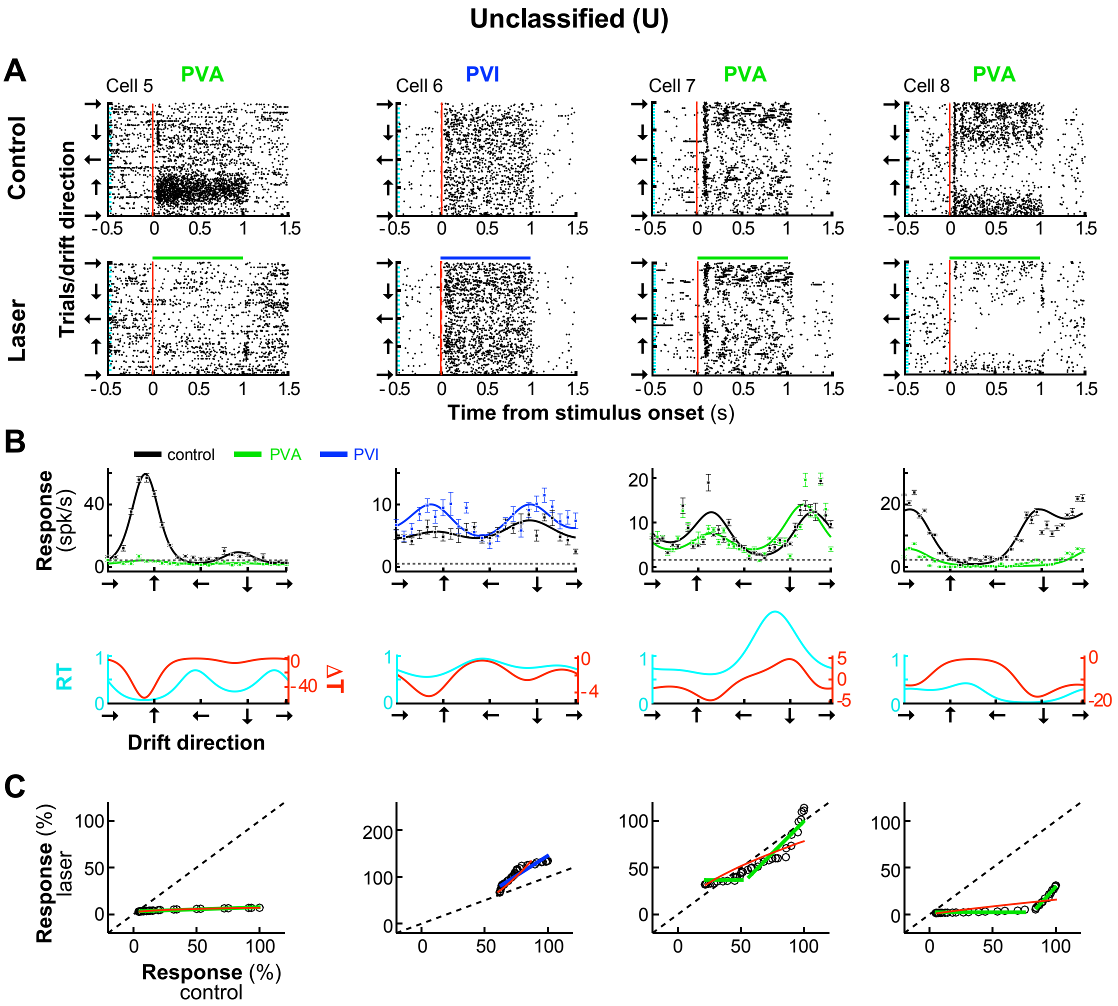


**Figure S3** (Related to Fig. 1)

**Unclassified units.**

Four example unclassified units. **(A)** Each column shows raster plots for an example Pyr unit in control (Top) and laser (PVA or PVI; Bottom) conditions. *Green and blue bars* at the top of the bottom rasters: time of laser photostimulation which was simultaneous with visual stimulus presentation (1s). Other conventions as in Fig. 1B. **(B)** Top: orientation tuning curves in control (*black*) and laser (PVA, *green;* PVI, *blue*) condition, fitted with a von-Mises function. Error bars: s.e.m. Bottom: $\Delta T$ curve (*red*) and $RT$ curve (*cyan*) for each respective cell. **(C)** Relative spiking response (% of max response in control condition) of the same units to stimuli of different drift directions in control vs laser conditions. *Green and blue lines*: threshold linear model fits. *Red line*: fits of our model described in Figs. 7-8.

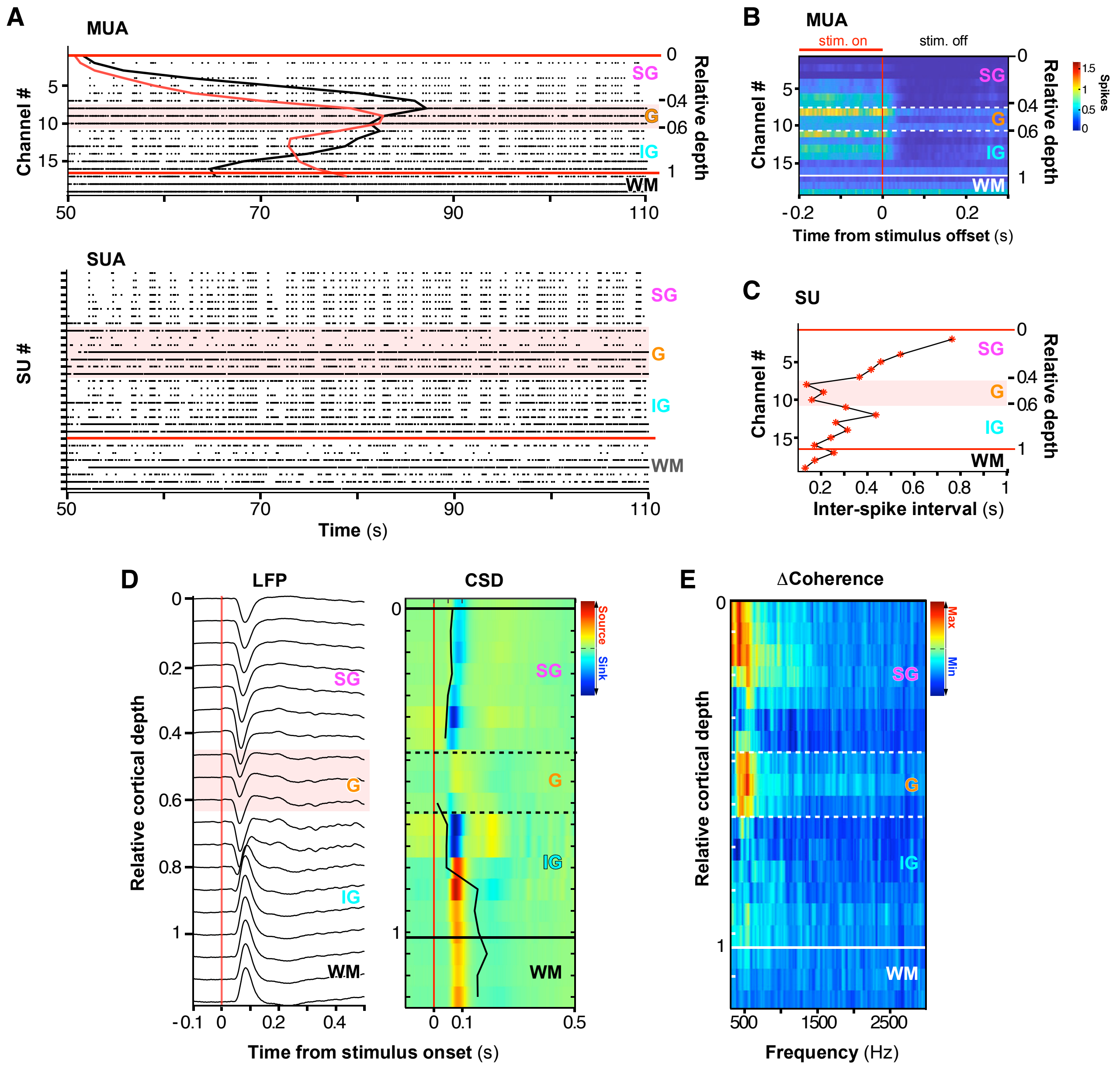


**Figure S4** (Related to Figs. 4-5)

**Identification of layer boundaries using laminar recordings.**

**(A)** Top: Raster plots of multi-unit activity (MUA) across LEA channels over a 60 sec period of recordings, for one example LEA penetration. The left y- axis indicates channel number (channel #1 is the first channel inside the cortex), while the right y-axis indicates the relative cortical depth, where 0 is the top of the cortex (corresponding to channel #1) and 1 is the border between layer 6 and the white matter (*WM*). The *red curve* indicates the mean spiking rate during the interstimulus intervals, and the *black curve* is the mean spiking rate over the entire recording periods (stimulus-on and interstimulus interval). *Red horizontal lines* mark the top and bottom of the cortex, and the *pink shading* highlights the granular (*G*) layer. *SG:* supragranular layers; *IG:* infragranular layers. Bottom: Raster plots of single unit activity (SUA) arranged by channel position, with multiple SUs recorded at the same channel; thus here the stack of recordings does not correspond to cortical depth. A higher firing rate is characteristic of the G layer. **(B)** Spike-triggered averages of MUA across the depth of the cortex around the time of stimulus offset (time 0). Elevated spontaneous activity after stimulus offset is typical of the G layer. *White dashed lines* demarcate the G layer boundaries; *solid black and white lines*: top and bottom of the cortex, respectively. **(C)** Mean inter-spike interval (ISI; for values >100 ms) for each channel. Channels 8–10 show lower average ISI values, indicative of higher firing rates. **(D)** Left: Stimulus-evoked LFP profile across layers for one example LEA penetration. *Red vertical line*: time of stimulus onset. Right: Baseline-corrected (z-scored; see STAR Methods) CSD calculated from the LFPs and displayed as a color map. *Black contour* indicates estimated onset latency of current sinks, used to determine the location of the earliest current sink, which identifies the G layer. *Solid black horizontal lines* indicate the top and bottom of the cortex; *dashed black lines:* G layer boundaries. **(E)** Stimulus-evoked local coherence spectrum (stimulus evoked-baseline). For each contact the coherence with adjacent contacts was estimated across frequencies (see STAR Methods); coherence is higher in the G layer (contacts 8-10).

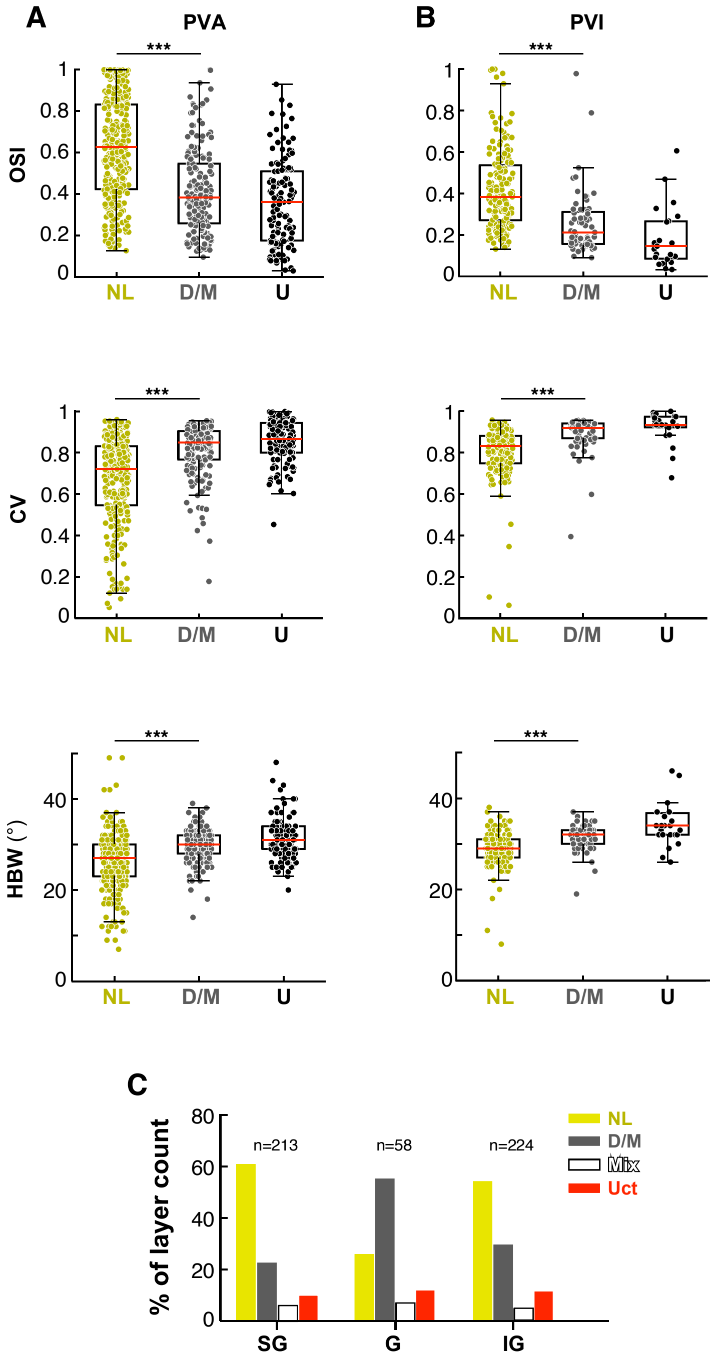


**Figure S5** (Related to Fig. 4)

**Tuning properties and laminar distribution of cells grouped by effect type.**
**(A)** Box plots of OSI (Top), CV (Middle) and HBW (Bottom) for NL (*yellow dots*), D/M (*gray dots*) and U (*black dots*) Pyr neurons recorded in the control (no laser) condition in PVA experiments. *Red horizontal lines*: median. **(B)** Same as in (A) but for units recorded in the control condition in PVI experiments. **(C)** Percent distribution of NL, D/M, and U (divided in Mix and Uct) units in different layers. n= total number of units in each layer.

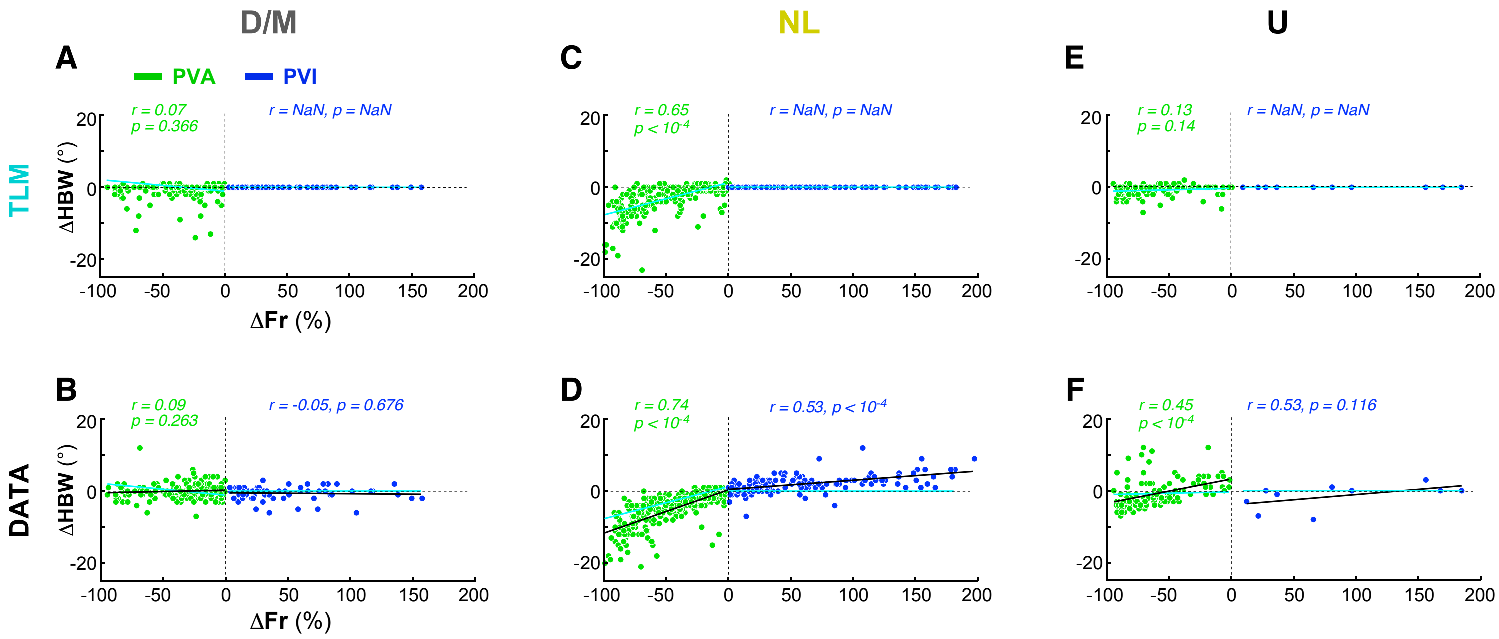


**Figure S6** (Related to Figs. 2, 8)

**The TLM does not capture the changes in HBW caused by $PV^+$ neuron manipulation in the data.**
**(A)** Scatter plot of the change in HBW as a function of the percent change in firing rate for TLM-simulated effects of $PV^+$ neuron manipulations on the D/M population. Here an in all remaining panels, the *cyan line* is the linear regression fit to the TLM-simulated data. The *r* and *p* values here and in panels (C, E) refer to the linear regression fit to the TLM-simulated data. **(B)** Same as in (A) but for the D/M population in the real data (same plot as shown in **Fig. 2A**). The *black line*, here and in (D,F) is the linear regression fit to the real data, to which the *r* and *p* values refer. **(C,D)** Same as (A,B) but for the NL population. Panel (D) shows the same plot as in **Fig. 2E**. **(E,F)** Same as in (A,B) but for the Unclassified (U) population. Other conventions are as in **Fig. 2A,E**.

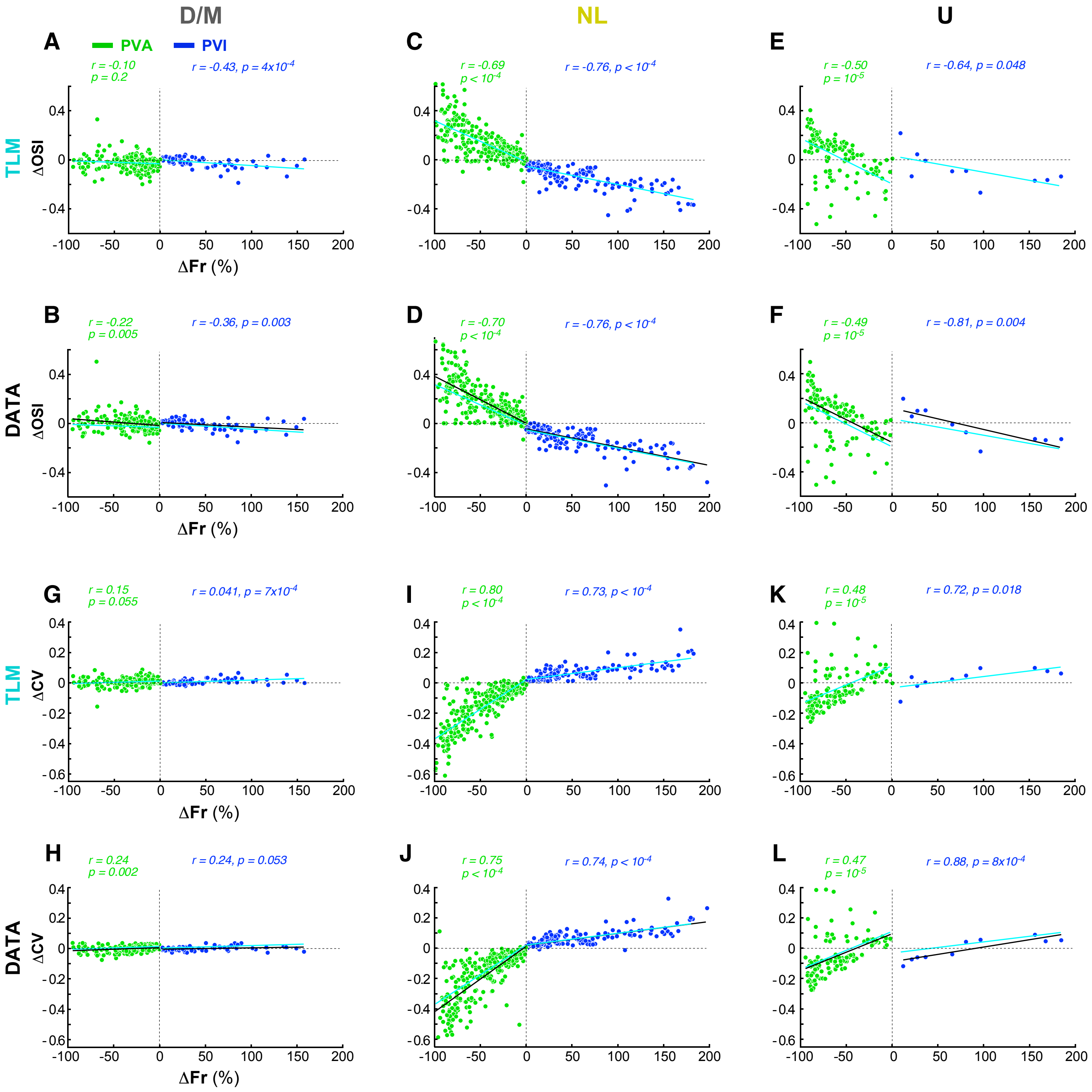


**Figure S7** (Related to Figs. 3,8)

**TLM-simulated changes in OSI and CV caused by $PV^+$ neuron manipulation.**
**(A)** Scatter plot of the change in OSI as a function of the percent change in firing rate for TLM-simulated effects of $PV^+$ neuron manipulations on the D/M population. Here an in all remaining panels, the *cyan line* is the linear regression fit to the TLM-simulated data. The *r* and *p* values here and in panels (C,E,G,I,K) refer to the linear regression fit to the TLM-simulated data. **(B)** Same as in (A) but for the D/M population in the real data (same plot as shown in **Fig. 3A**). The *black line*, here and in (D,F,H,J,L) is the linear regression fit to the real data, to which the *r* and *p* values refer. **(C,D)** Same as (A,B) but for the NL population. Panel (D) shows the same plot as in **Fig. 3C**. **(E,F)** Same as in (A,B) but for the Unclassified (U) population. Other conventions are as in **Fig. 3A,C**. **(G,H)** Same as in (A,B) but here for the D/M population, we show the TLM-simulated (G) and real (H) changes in CV as a function of change in firing rate. Panel (H) shows the same plot as in **Fig. 3E**. **(I,J)** Same as in (G,H) for the NL population. Panel (J) shows the same plot as in **Fig. 3G**. **(K,L)** Same as in (G,H) but for the U population.

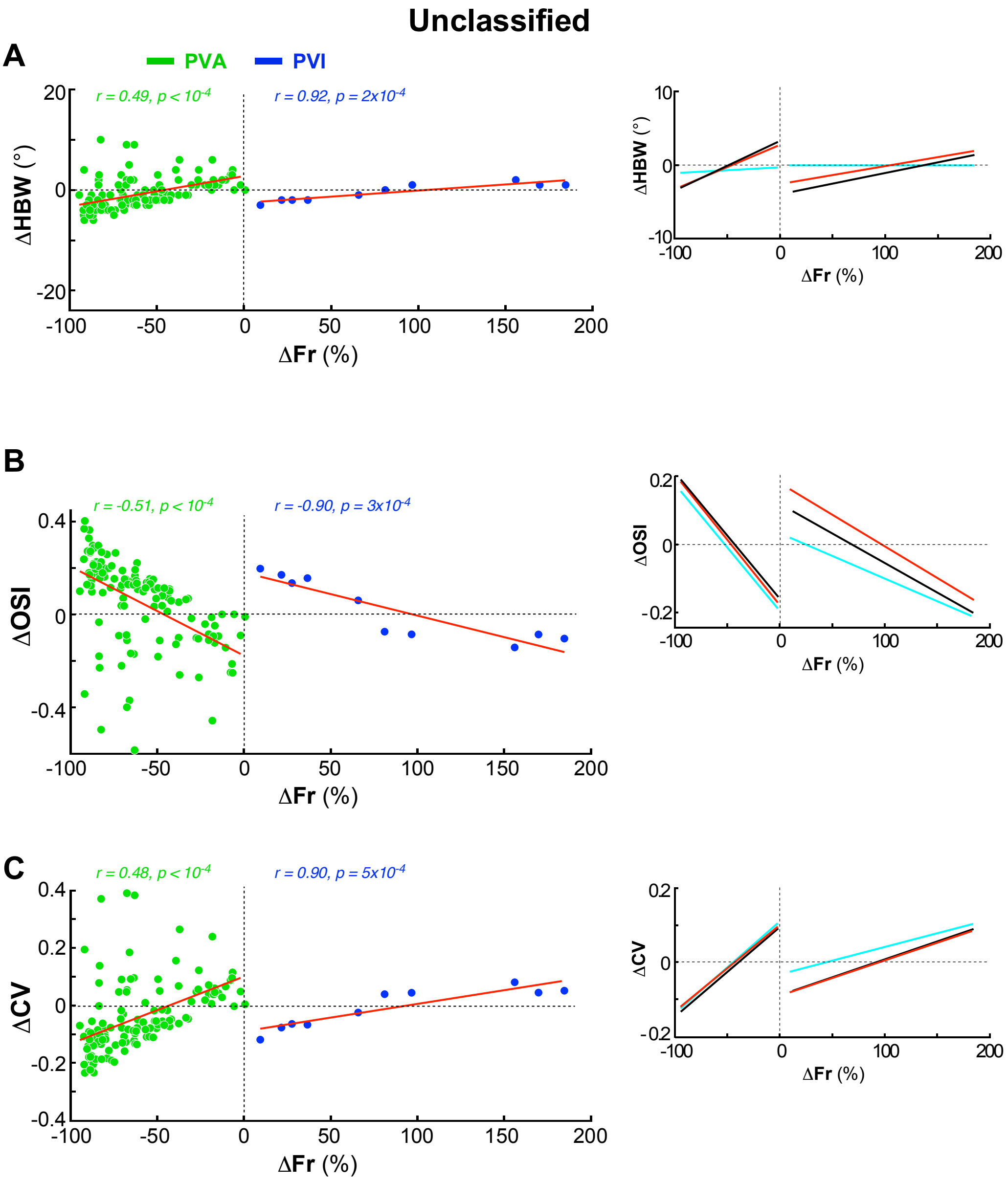


**Figure S8** (Related to Fig. 8)

**I/O model-simulated changes in HBW, OSI and CV caused by $PV^+$ neuron manipulation on the Unclassified cell population.**

**(A)** Scatter plot of the change in HBW as a function of the percent change in firing rate for I/O model-simulated effects of $PV^+$ neuron manipulations on the Unclassified population. Here and in all remaining panels, the *red line* is the linear regression fit to the simulated data, to which the *r* and *p* values refer. *Right inset*: comparison of linear regression fits to the data (*black line*), the TLM-simulated data (*cyan line*) and the I/O model-simulated data (*red line*). **(B)** Same as in (A) but for changes in OSI. **(C)** Same as in (A) but for changes in CV. Other conventions are as in **Figs. 2-3**.
